# Single molecule-level detection and long read-based phasing of epigenetic variations in bacterial methylomes

**DOI:** 10.1101/007823

**Authors:** John Beaulaurier, Shijia Zhu, Robert Sebra, Xue-Song Zhang, Chaggai Rosenbluh, Gintaras Deikus, Nan Shen, Diana Munera, Matthew K. Waldor, Martin Blaser, Andrew Chess, Eric E. Schadt, Gang Fang

## Abstract

Comprehensive genome-wide analyses of bacterial DNA methylation have not been possible until the recent advent of single molecule, real-time (SMRT) sequencing. This technology enables the direct detection of N6-methyladenine (6mA) and 4-methylcytosine (4mC) at single nucleotide resolution on a genome-wide scale. The distributions of these two major types of DNA methylation, along with 5-methylcytosine (5mC), comprise the bacterial methylome, some rare exceptions notwithstanding. SMRT sequencing has already revealed marked diversity in bacterial methylomes as well as the existence of heterogeneity of methylation in cells in single bacterial colonies, where such ‘epigenetic’ variation can enable bacterial populations to rapidly adapt to changing conditions. However, current methods for studying bacterial methylomes using SMRT sequencing mainly rely on population-level summaries that do not provide the single-cell resolution necessary for dissecting the epigenetic heterogeneity in bacterial populations. Here, we present a novel SMRT sequencing-based framework, consisting of two complementary methods, for single molecule-level detection of DNA methylation and assessment of methyltransferase activity through single molecule-level long read-based epigenetic phasing. Using seven bacterial strains and integrating data from SMRT and Illumina sequencing, we show that our method yields significantly improved resolution compared to existing population-level methods, and reveals several distinct types of epigenetic heterogeneity. Our approach enables new investigations of the complex architecture and dynamics of bacterial methylomes and provides a powerful new tool for the study of bacterial epigenetic control.

## Introduction

In the bacterial kingdom, DNA methylation is catalyzed by three families of DNA methyltransferases (MTases) that typically add methyl groups to DNA bases in a sequence-specific manner^1–3^. One family of MTases attaches methyl groups to adenine residues, creating N6-methyladenine (6mA), whereas the other two families introduce methyl groups to cytosine residues to create either N4-methylcytosine (4mC) or 5-methylcytosine (5mC). Many bacterial DNA MTases act in concert with and are encoded in close proximity to cognate restriction endonucleases (REs); the MTase protects DNA from digestion by the RE with which it forms a restriction-modification (RM) system^2,4,5^.RM systems are generally believed to have an ‘immune’ function, protecting cells from invading foreign DNA. They are also studied as selfish elements that protect themselves against removal through the post-segregational killing of new progeny by pre-existing and stable RE molecules^4, 6,7^.In addition, so-called ‘orphan’ MTases, which occur in the genome without an associated RE, have been found to play important regulatory roles in global gene expression and other biological processes^2,3,8–10^.Furthermore, the ability of such MTases to target their recognition motifs for methylation often depends on competitive binding at the target site between several DNA binding proteins^2,3,1^1–15. These epigenetic regulators of gene expression, including both MTases and competing DNA binding proteins, are a source of *phase variation*^2,3,16^ that increases the robustness of the population and provides opportunities to adapt in response to changing environmental conditions^17–19^.

In some bacteria, the behavior and role of certain MTases can vary dramatically due to slippage events during DNA replication through homopolymer-rich MTase coding sequence. This slippage can cause frameshift mutations that result in truncated and usually inactive MTases^20–22^ or in active MTases with altered target sequence specificity^23,24^. These mechanisms can yield heterogeneity in the methylomes present within descendants of a single population and can cause differential regulation of multiple genes, termed phase-variable regulon (*phasevarion)*^20–22^. It has been postulated that epigenetic control of gene expression mediated by phase variation (epigenetic control of a single gene) or phasevarions (multiple genes regulated simultaneously) allows a given essentially clonal population to adopt multiple distinct phenotypes. Such heterogeneity can facilitate adaptation to diverse environmental niches, including complex host environments and presence of antibiotics, as has been reported in several studies^22,25–27^.

Genomic analyses suggest that some form of DNA methylation is present in nearly all bacteria, as putative DNA MTases were found in 94% of 3300+ sequenced bacterial genomes^5^. Given the large number of MTase target sites in bacterial genomes and the growing evidence suggesting regulatory roles of methylation by both RM and orphan MTases^28,29^, the potential scope for exploring the diversity of bacterial methylation and methylation-mediated gene regulation is vast. Previous methods have been laborious, often lacking single nucleotide resolution and whole genome scalability, but have nevertheless demonstrated the critical roles that epigenetic variation plays in a great diversity of processes^2,3,11,13–15^. However, the precise sequence targets and biological roles of most MTases, including their dynamics and functions, remain virtually unknown. While recent progress in bisulfite sequencing facilitates the accurate detection of 5mC methylation and has been successful in studying 5mC sites in *E. coli*^30^, there has been a lack of a high throughput genome-wide sequencing methodology for efficiently detecting 6mA and 4mC methylation in bacteria.

Single molecule, real-time (SMRT) DNA sequencing technology^31^ represents a major advance with respect to the detection of nearly twenty different types of chemical modifications to DNA, including all three major types of DNA methylation in bacteria (6mA, 4mC, and 5mC), although the reduced signal-to-noise ratio with 5mC makes detection of such events with SMRT sequencing a challenge. Through SMRT sequencing, each DNA molecule, consisting of both strands of native DNA fragments circularized by ligating hairpin adapters to the ends, is sequenced by a DNA polymerase, where due to the circular topology a given fragment can be sequenced multiple times in the real-time DNA synthesis process. The SMRT sequencing instrument not only monitors the pulse fluorescence associated with each incorporated nucleotide, but also records the time between the incorporation events, termed the inter-pulse duration (IPD). Variation in IPDs (referred to as kinetic variation) is highly correlated with the presence of modifications within the DNA template^32–34^, including 6mA, 4mC, 8-Oxoguanine etc.

SMRT sequencing has been applied to genome-wide characterization of several bacteria^34–36^. These genome-wide studies, using motif enrichment analyses, enabled the comprehensive *de novo* identification of specific recognition sites targeted by the MTases encoded in these genomes. The application of SMRT sequencing to the characterization of a growing number of bacterial methylomes^23,24,37,38^ has revealed both numerous MTases with novel specificities and great diversity in methylomes among bacterial species and strains^15,27,28,44^. However, while the accurate identification of methylation motifs has enabled these discoveries, the heterogeneity and dynamics of methylomes within cultured bacterial populations and under different conditions have not been thoroughly explored. This is in large part due to limitations of current SMRT sequencing-based bacterial methylome analysis protocols, which rely mainly on assessments of aggregate IPD values at each genomic position across populations of cells^32–35,39.^ With this approach, the individual reads (corresponding to molecules) are aligned to the same genomic region and then statistically modeled as an ensemble (Fig. 1a, b). This approach enhances the statistical power for methylation detection at single nucleotide resolution and improves motif-level analysis, but fundamentally limits the ability to resolve the subtle dynamics and heterogeneity in bacterial methylomes. Although some progress in the investigation of dimensions beyond traditional population-level analysis have recently been reported^37,40^, new methods are needed to decipher the heterogeneity and dynamics of bacterial DNA methylation at single molecule resolution.

**Figure 1.**
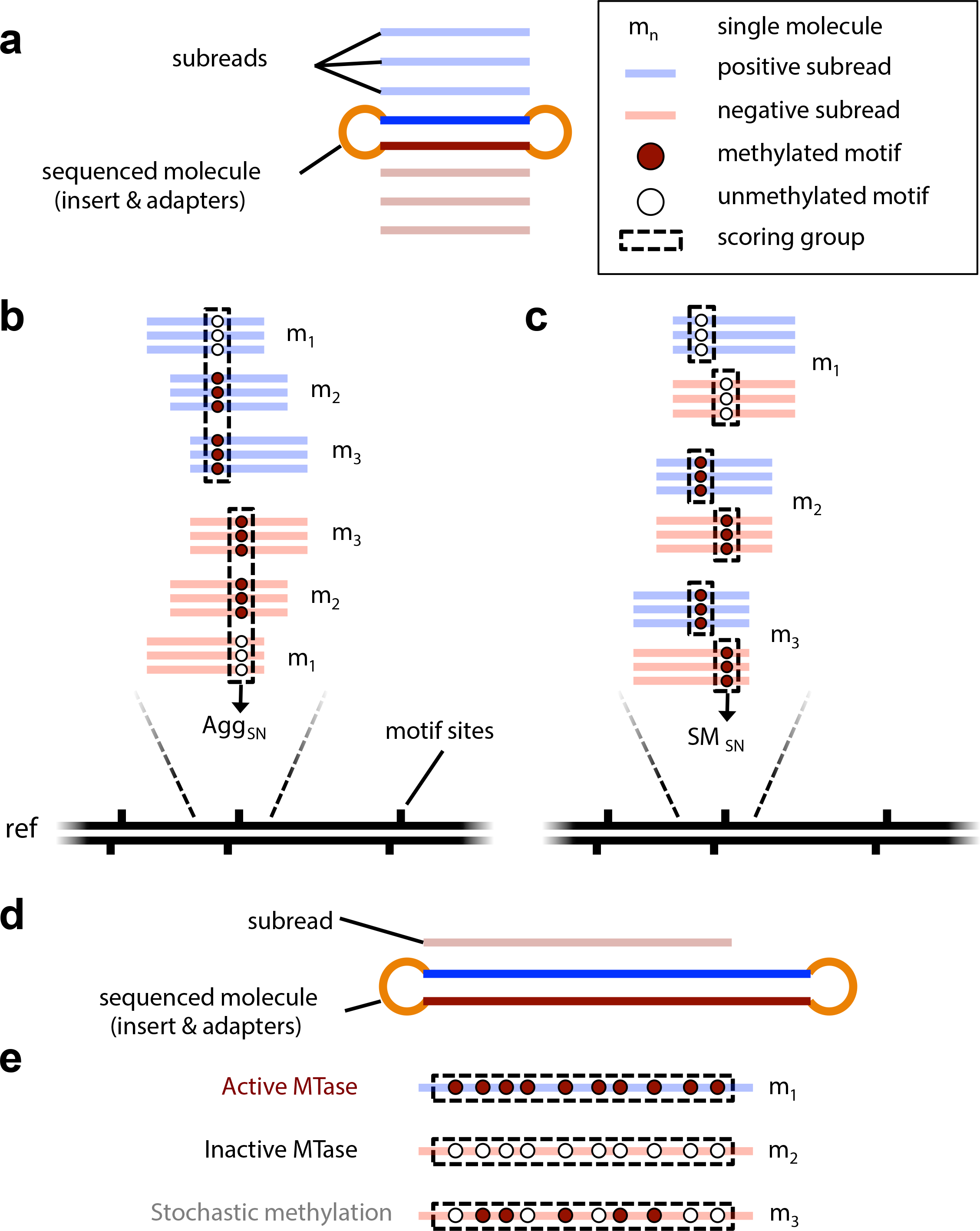
Schematic illustrating the general approaches of both the existing and two proposed methods for detecting DNA methylation in SMRT sequencing reads. **(a)** Illustration of an individual SMRT sequencing molecule (short DNA insert + adapters) used for the single molecule, single nucleotide (SM_sn_) detection method and the subreads that are produced during the sequencing process. **(b)** The existing methylation detection method relies on an aggregated treatment of individually sequenced molecules. For a given strand at each genomic position, the IPD values from all the subreads aligning to that strand and position are aggregated together to infer the presence of a methylated base. **(c)** The proposed SM_sn_ method for detecting DNA methylation. Instead of aggregating molecules together to infer a single methylation call for each genomic position and strand, the methylation calls for each genomic position and strand are made at the level of each molecule using only the IPD values from its own subreads to make a molecule-specific determination. **(d)** Illustration of an individual SMRT sequencing molecule (long DNA insert + adapters) used for the single molecule pooled (SM_p_) and a typical long subread that is produced during the sequencing process. **(e)** The proposed SM_p_ approach for measuring MTase activity and detecting possible phasevarions. This approach retains the single molecule resolution, but instead of generating methylation scores for each individual motif site on the molecule, it pools together IPD values from multiple distinct motif sites along the length of the long read to assess MTase activity on the sequenced molecule.

Here, we present a novel and systematic framework for single molecule-level detection of bacterial DNA methylation and assessment of MTase activity using SMRT sequencing data. The foundation of this approach rests on two complementary methods that use molecule-specific IPD information to infer methylation states at single molecule resolution (Fig. 1c, e). We demonstrate the framework’s effectiveness through comprehensive, quantitative characterization of seven bacterial methylomes and by identifying several types of heterogeneity in these methylomes. The enhanced resolution provided by our analytic framework broadens our knowledge of bacterial methylomes and enables deeper understanding of the diverse roles of methylation in modulating bacterial physiology.

## Results

### Methods for single molecule-level detection of methylation and epigenetic phasing

We first present the rationale and description of the two complementary methods for (i) single molecule, single nucleotide, strand-specific detection of methylation events and (ii) single molecule-level epigenetic phasing analysis. Our strategy revolves around the interrogation of circular consensus sequence data generated from short-insert SMRT sequencing libraries and continuous long read data generated from long-insert SMRT sequencing libraries^31,41^. For short insert libraries, DNA molecules are cleaved into ∼250bp double-stranded fragments and hairpin adaptors are ligated to each end. The sequencing-by-synthesis occurs when the DNA polymerase begins synthesizing DNA using the short fragment as a template. Since the average read length in the SMRT sequencing system is 8,500bp, this same short DNA fragment can be sequenced repeatedly by the same DNA polymerase (each strand can be read > 15 times on average). The multiple passes over the same molecule (Fig. 1a-c) allow the calculation of a single molecule, single nucleotide (SM_sn_) score and provide statistical power to detect shifts in the kinetic variation at a specific nucleotide position on a given DNA molecule.

Complementing the short insert libraries are the long insert libraries, which given the read length of the SMRT sequencing system, enables sequencing of long stretches of DNA. This long library sequencing has been successfully applied in *de novo* bacterial genome assembly for its unique ability to resolve genomic regions with complex repeated elements^41–45^. Here, we instead leverage the continuous long reads for methylation phasing at single molecule resolution. For a particular methyltransferase of interest, its target sequence motif may be represented multiple times in a given long stretch of DNA. For example, if the target motif is 4 nucleotides long, then that motif will be represented 33 times on average in a strand of DNA 8,500bp long. The kinetic variation statistics for each occurrence of a given motif on a single strand of DNA can then be pooled to increase statistical power to infer whether a given motif is being actively methylated in a particular bacterial cell (Fig. 1d, e). Since methyltransferases often methylate almost all occurrences of their target sequence, this single molecule pooled (SM_p_) score can provide strong evidence as to whether a given methyltransferase is active in a bacterial cell. Furthermore, the precise value of SM_p_ can also be used to infer the processivity of a methyltransferase of interest, as is discussed later. At current average SMRTseq read lengths (∼8,500bp), this estimation procedure will be limited to shorter methylation motifs for most of the molecules, as longer motifs (e.g., > 5mers) occur less frequently. However, the very longest reads (read lengths up to 60,000bp are now possible) provide an opportunity to estimate the SM_p_ scores accurately for longer motifs.

Therefore, the short-library based SM_sn_ scores can be used to estimate the methylation state of a particular occurrence of a given motif on a single molecule, while the long-library based SM_p_ scores enable the assessment of the activity of a methyltransferase targeting a given motif through single molecule-level epigenetic phasing. Details of the two methods can be found in Online Methods. Next, we evaluate the performance of the two methods, and demonstrate how SM_sn_ and SM_p_ scores can be used to characterize distinct types of epigenetic heterogeneity in different bacteria.

### Sensitivity and specificity of single-molecule, single nucleotide (SM_sn_) detections

Confident estimation of the IPD ratio relies on an accurate mean IPD for each nucleotide in a single molecule. The accuracy of the mean IPDs increases as the number of subreads for each molecule increases^32,33^, since each subread provides an independent estimate of the IPD. With an average read length (currently ∼8,500bp), the number of subreads (i.e. single molecule coverage, cov_sm_) is negatively correlated with the insert size of the sequencing library. To quantify the sensitivity and specificity for single molecule, single nucleotide-level detection, we tested this framework on methylated 5’-CTGC<u>A</u>G-3’ sites in a native *E. coli* O104:H4 C227-11strain^34^ (C227), and a matching whole genome amplified (WGA) sample in which the methylation sites were erased (Online Methods; Fig. 2a). As expected, the sensitivity of the method increases as cov_sm_ increases. For the single molecule, single nucleotide approach, when cov_sm_ ≥ 15 (corresponding to an average library size of 280bp), 6mA can be detected with a sensitivity of 98.5% and a specificity of 99.5%. This suggests that 6mA can be accurately detected at a single molecule resolution using a short insert library. Although a smaller library size can provide higher values of cov_sm_, a minimum library size of 150-200bp is recommended to avoid loss of genomic DNA during library construction and to facilitate removal of adapter-dimer constructs during purification.

**Figure 2.**
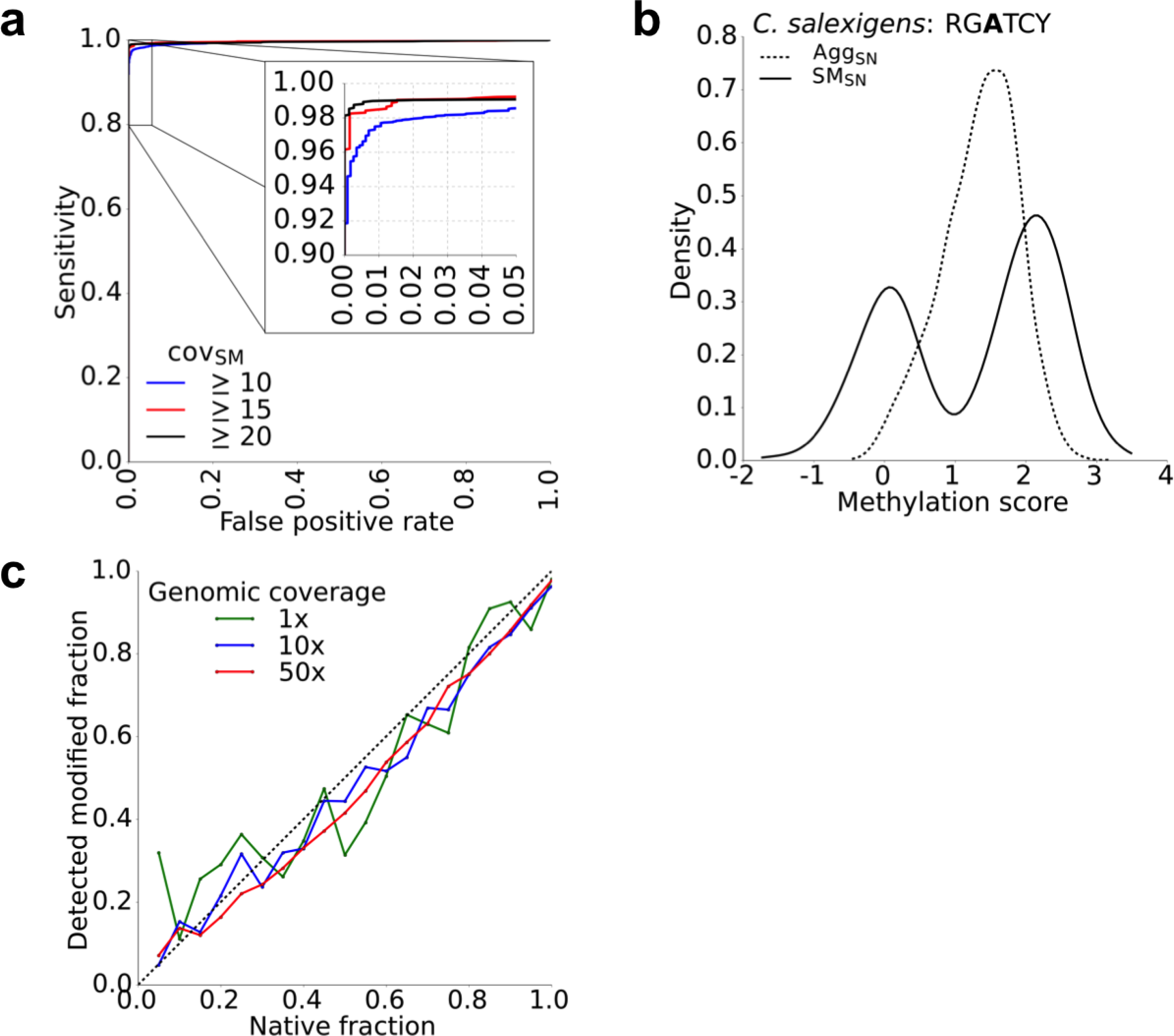
Multiple metrics showing the performance of the proposed single molecule, single nucleotide detection method. **(a)** Performance of the approach for detecting 6mA modifications in the 5’-CTGCAG-3’ motif of *E. coli* C227 using three thresholds for minimum single-molecule coverage (cov_sm_). **(b)** Probability density illustrating the aggregate, single nucleotide (Agg_SN_) and single molecule, single nucleotide (SM_sn_) methylation scores for 5’-RGATCY-3’ in *C. salexigens,* a motif which is known to have a large non-methylated fraction. The bimodal distribution provided by the SM_sn_ scores enables the accurate and objective estimation of this fraction. **(c)** Accuracy of SM_sn_-enabled estimations of the methylated fraction (using cov_sm_≥10) for the 5’-CTGCAG-3’ motif of *E. coli* C227 at various levels of genomic sequencing coverage.

We also estimated the sensitivity and specificity of detecting 6mA at single molecule resolution for other 6mA motifs and found the results to be comparable to CTGCAG (Supplementary Fig. 1a, b). Upon interrogating a 4mC motif, we found that 4mC can also be accurately detected at single molecule resolution (Supplementary Fig. 1c). For 5mC, we did not attempt similar estimation at single molecule resolution because of a much lower signal-to-noise ratio, even after conversion of 5mC to 5hmC with Tet enzymes^46^, consistent with other recent observations ^23,37.^ The remainder of this study will focus solely on the characterization of 6mA, the most prevalent type of methylation in bacteria, although all of the analyses detailed are applicable to 4mC in addition to 6mA.

### Distribution of SM_sn_ scores enhance quantification of global methylation heterogeneity

The methylome of *Chromohalobacter salexigens* was characterized in a recent study and found to contain a substantial number of non-methylated 5’-RGATCY-3’ sites^36^. Specifically, 23.5% of the motif sites were predicted to be non-methylated based on the standard population level analysis depicted in Figure 2b (Online Methods), consistent with the original study^36^. As shown, the distribution of molecule-aggregated single nucleotide (Agg_SN_) scores does not show clear separation between methylated and non-methylated RGATCY sites, indicating that the estimated percentage of methylation depends on a subjective, ad hoc threshold. In contrast, clear bimodality is observed in the distribution of SM_sn_ scores, where the individual components centered near SM_sn_≈0 and SM_sn_≈2 represent the non-methylated and methylated fractions, respectively (Fig. 2b). Note that, a distribution of SM_sn_ centered at zero correspond to motif sites that are not methylated because the IPDs do not differ between native and WGA DNAs; in contrast, a distribution of SM_sn_ centered near 2 corresponds to motif sites that are methylated (SM_sn_≈2 corresponds to an IPD ratio of ∼7 in log scale, specific to the P4 sequencing chemistry and 6mA; Supplementary Figs. 3–8). This bimodality allows the percentage of methylated motif sites to be objectively estimated at 60.4% using a standard expectation maximization (EM) algorithm^47^ (Online Methods) without the need for a subjective input threshold.

**SM_sn_-based quantification is stable at low sequencing coverage:** Next, we designed an *in silico* experiment to evaluate the reliability of the methylated percentage estimation based on the SM_sn_ score distributions. To generate *in silico* methylated fractions, we mixed single sequencing molecules from the native *E. coli* C227 data and corresponding WGA molecules, starting at 5% native molecules and increasing the native fraction stepwise by 5% until we reached 100% native molecules. The percentage of methylated and non-methylated sites in each *in silico* mixture was estimated using the distribution of SM_sn_ scores for CTGCAG sites (cov_sm_≥10). Because we expect most CTGCAG sites in the native molecules to be methylated, an accurate estimation is one in which the detected methylated fraction closely approximates the known native fraction in each mixture. Additionally, because the estimation of methylated fractions could provide added value into the characterization of *in vivo* isolates where sequencing coverage is often an issue, we evaluated the accuracy of our methylated fraction estimations at varying levels of genomic sequencing coverage. By down-sampling the data (Online Methods), we found that the methylated fraction estimations are highly stable even when global coverage is 1x (Fig. 2c). This result supports the applicability of this approach to *in vivo* isolates from which very limited amounts of native DNA are available, leaving low coverage SMRTseq sequencing as the only option. We considered the possibility that the systematic downward shift from the diagonal in the estimated methylated fraction could be due to a bias in the EM algorithm or the assumption that SM_sn_ follows a normal distribution, causing a skew in the fraction estimates. However, two independent tests using mixtures of simulated distributions did not support this hypothesis (Supplementary Fig. 2; Online Methods). This suggests that the downward shift likely reflects that a small number of the CTGCAG sites in the native molecules are non-methylated, causing a slight underestimation of the fraction of native molecules.

**Global methylation heterogeneity in six bacteria:** We applied the SM_sn_ based quantification analysis to six bacterial methylomes that were recently sequenced^34,36,37^ or specifically sequenced for this study (Supplementary Table 1). We first detected methylation motifs based on existing methods^23,34,36^ and divided them into two groups, based on the global distribution of SM_sn_ scores. In the first group, the majority of motif sites (> 95%) were methylated, while only a very small proportion were non-methylated, likely due to competitive binding between the MTases and other DNA binding proteins such as transcription factors^2,3,11,13,14^. In the second group, a substantial percentage of motif sites were non-methylated (> 5%), which may suggest the existence of alternative mechanisms that drive the heterogeneity of the methylome. Some representative bacterium-motif pairs belonging to each of the two groups were shown in Figures 3b and 3c, respectively. A complete SM_sn_ score summary for each bacterium is shown in Supplementary Figures 3–8. While most motifs belong to the first group and do not show extensive non-methylation, the second group includes the RGATCY motif of *C. salexigens* and three motifs from *Helicobacter pylori* J99. Most intriguingly, the *H. pylori* motif 5’-GWC<u>A</u>Y −3’ shows an extremely high percentage of non-methylated sites (75.3%), which we will investigate in-depth using long library based phasing analysis.

**Figure 3.**
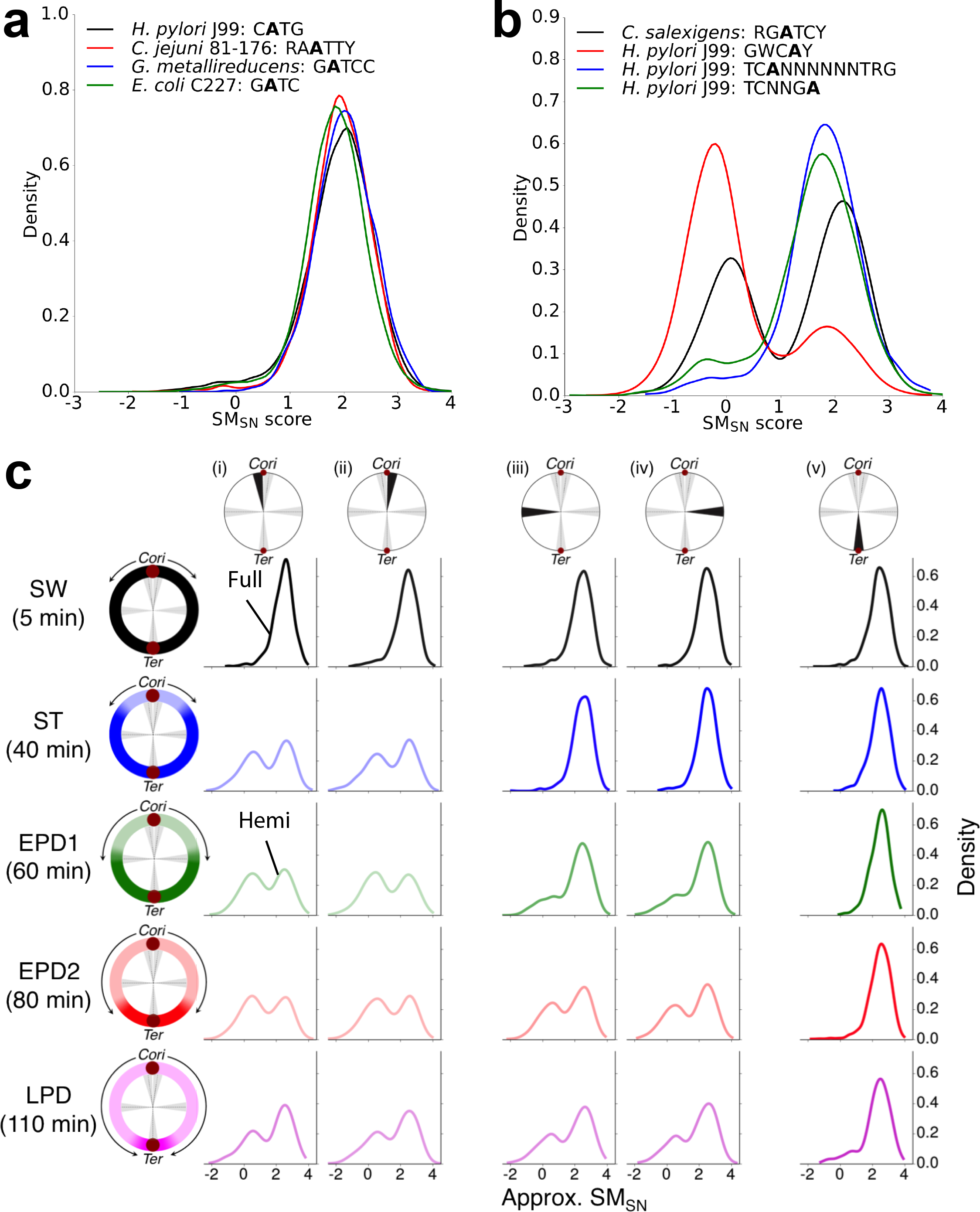
Examples of various levels of heterogeneity in multiple bacterial methylomes as observed using single molecule, single nucleotide detection. **(a)** SM_sn_ score distributions for multiple bacterium-motif pairs that exhibit near complete methylation, along with a non-methylated motif for comparison. These examples show little or no bimodality, but are rather almost entirely distributed around SM_sn_ ∼ 2, indicating that the vast majority of motif sites are methylated. **(b)** SM_sn_ score distributions for multiple bacterium-motif pairs that display significant non-methylated fractions. The bimodality of each distribution enables an accurate estimation of the methylated and non-methylated fractions. The *H. pylori* J99 motifs show minor variation in the SM_sn_ associated with each peak due to subtle differences in the chemistry version used for SMRT sequencing of their native and WGA samples. **(c)** Single molecule, single nucleotide level interrogation of GANTC methylation at five genomic positions (columns) in synchronized *C. crescentus* during a single round of DNA replication. Five time points (rows) provide snapshots of the bidirectional progression of the replication forks as they proceed in both directions away from the origin of replication *(Cori)* to the terminus *(Ter)* on the *C. crescentus* chromosome. Gray wedges in the chromosome schematics show the 200-kilobase genomic windows that are interrogated. Two regions are on either side of the *Cori:* (i) *Cori* −0.1 megabase and (ii) *Cori* + 0.1 megabase. Another two are halfway between *Cori* and *Ter:* (iii) *Cori* −1 megabase and (iv) *Cori* + 1 megabase. The final region covers the terminus: (v) *Ter.* For each time point, light (hemimethylated) to dark (fully methylated) color shading in the schematic and PDF curves illustrate the approximate position of the replication fork. Progressive hemi-methylation of GANTC sites follows the replication fork and is apparent by the bimodal distribution of the approximate SM_sn_ scores (approximate due to lack of WGA sequencing, see Online Methods). Hemimethylated sites cannot transition back to full methylation until transcription of the gene encoding the proper methyltransferase, CcrM, which does not occur until very late in the replication process. The late pre-divisional (LPD) time point captures some of this transition, as stochastic variation in the replication speed means that some of the replication rounds have likely finished and are transcribing CcrM.

### Distribution of SM_sn_ scores can enhance interpretation of regional methylation dynamics

The *Caulobacter crescentus* genome encodes a methyltransferase (CcrM) targeting 5’-G<u>A</u>NTC-3’ sites. The corresponding gene, ccrM, is only expressed at the late stage of the cell cycle^48^. Consequently, fully methylated GANTC sites transition to hemi-methylated as the replication fork proceeds from the origin of replication (*Cori*) to the terminus *(Ter)*^37^. Recently, Kozdon et al.^37^ used SMRT sequencing to study such transitions at five time points over the cell cycle. This provides a good dataset for testing whether SM_sn_ distributions can be used to quantitatively interpret methylation dynamics of specific genomic regions. We sampled the genome of *C. crescentus* at five distinct 200-kilobase regions at five different time points during a single synchronized round of cell division (Fig. 3c). As the replication fork proceeds from the *Cori* to the *Ter* in the five time points, the SM_sn_ score distributions reveal an increasing fraction of the genome that has been converted from fully methylated to hemi-methylated GANTC sites. At the first time point, the fully methylated state of the swarmer cells (SW) is apparent by the single mode in SM_sn_ scores at all five genomic regions. As the cells differentiate into the actively replicating stalked cells (ST), there are bimodal SM_sn_ scores in regions (i) and (ii) that are closest to the *Cori,* reflecting the passage of the replication fork through those regions. The following two time points, early pre-divisional stages (EDP1 and EDP2, respectively), reveal the transition of the single-mode SM_sn_ distributions in regions (iii) and (iv) to clear bimodal distributions, reflecting the passage of the replication fork through those regions. The unequal bimodal distribution in EDP2 in regions (iii) and (iv) indicates that the molecules from that region have not all been converted from fully- to hemi-methylated, likely due to stochastic differences in the position of the replication fork 80 minutes post-synchronization. The final time point, late pre-divisional (LPD), reveals a genome that has almost completely converted to a hemi-methylated state, with the exception of the region immediately surrounding the *Ter* (v), which only shows a small amount of hemi-methylation. In fact, regions (i)-(iv) at this final time point indicate that these sites have already begun to transition from equal bimodality (universal hemi-methylation) back toward their starting single-mode distributions (universal full methylation). This is also likely due to imperfect synchronization, allowing some replication forks to progress past the *ccrM* gene at the LPD time point, thus activating transcription and beginning the process of re-methylating all the GANTC sites that had been hemi-methylated by the passage of the replication fork.

### Distinct types of global epigenetic heterogeneity characterized by single molecule-level assessment of MTase activity

We envisioned three potential alternative explanations for the heterogeneous methylation of motifs that were frequently non-methylated (Fig. 3b). In the first, we hypothesized that the methylation motif may be inaccurate and the observed heterogeneity arises from a mixture of truly highly methylated motifs and motifs falsely identified by the motif enrichment algorithm. Alternatively, the MTase might stochastically methylate only a fraction of its recognition motif sites, in which case the fractions of methylation estimated based on global distribution of SM_sn_ scores reflect a universally active MTase, albeit one without the ability to methylate all of its target motifs. Finally, sequence variation in the MTase may cause a phasevarion. In this third case, mutations in homopolymers or other simple sequence repeats present in coding or regulatory sequences can cause switching that either activates/deactivates the MTase or switches its target specificity^20–24^. Either of these two switching modes can induce cell-wide methylation patterns that might differ within a single population. In this case, the fraction of methylation estimated by SM_sn_ scores reflects the fraction of cells with an active MTase targeting the methylation motif.

We can reject the first hypothesis because the SM_sn_ score distributions of explicit specifications of each motif showed similar heterogeneity (Supplementary Fig. 9), suggesting that the heterogeneity is not due to the mixing of true and false motifs. We can differentiate between the other two hypotheses by pooling SM_sn_ scores from distinct motif sites on an individual molecule as a way of phasing epigenetic information across the full length of the sequencing read (i.e. co-occurrence on a single molecule). If some molecules within a sample are methylated at all sites, while others are completely non-methylated, this supports the existence of a phasevarion. Because the expected frequency of methylation motif occurrence is often low (e.g. 1 motif per 256bp expected for a 4-mer motif, 1 motif per 1024bp expected for a 5-mer motif, etc.), the SMRT sequencing data used in the above analyses with short library sizes (250∼1000bp on average) provides limited statistical power to differentiate between the two hypotheses. The short library preparation generates molecules for sequencing that simply do not contain sufficient distinct motif sites to effectively determine whether or not a molecule is methylated via pooling its SM_sn_ scores. This motivated the design of the long library based SMp analysis for more effectively epigenetic phasing as described earlier (Fig. 1d, e).

***Direct observation of phasevarion and causal phase-variable MTases in H. pylori***. Phase-variable genes, including several MTases, are well-described in *H. pylori*^22,24,49,50^. In an effort to better understand the contributions made to the *H. pylori* methylome by phase-variable MTases, researchers have recently focused on identifying their targeted methylation motifs^23^. In that work, correction of inactivating frameshift mutations in several putative MTase genes was followed by SMRT sequencing of the resulting clonal populations. By generating modified and constitutively active versions of the putative phase-variable MTases, Krebes et al were able to assign methylation target motifs to the MTases using the existing, aggregation-based method for methylation detection. However, because most *H. pylori* isolates are likely not clonal and significant variation in homopolymer length has been observed between closely-related strains of *H. pylori*^49^, it is possible that active copies of the phase-variable MTases are already present in the isolates, just at levels too low to detect using the aggregation-based method. This provided an opportunity to evaluate the ability of the SMP-based method to detect minor subpopulations of active and inactive MTases.

We first tested the SM_p_ method on an *H. pylori* J99 isolate that was sequenced using libraries with long (∼20kb) DNA inserts. We targeted the 4-mer GATC motif (>95% of these sites are estimated to be methylated (Fig. 3a)) and tested the MTase activity for each sequenced molecule containing at least 10 motif sites. The distribution of SM_p_ scores, as well as a control distribution of SM_p_ scores where the IPD values have been randomly shuffled *in silico* between molecules, is shown in Figure 4a. No bimodality is present in the SM_p_ score distribution and it is nearly identical to the IPD-shuffled SM_p_ score distribution, suggesting that the MTase responsible for targeting the GATC motif in *H. pylori* J99 (M.Hpy99VI) is constitutively active. Through false discovery rate estimation (Methods), we found that only 0.07% of the molecules with at least 10 GATC sites from the *H. pylori* J99 long library data had evidence of non-methylation (maximum FDR = 1%). This level of non-methylation is consistent with several other motifs for which we expect to observe near-universal methylation activity (CATG, GANTC, and GAGG), suggesting that the small number of non-methylated reads may have originated from transiently hemi-methylated regions directly behind the DNA replication fork. Furthermore, phase-variation of M.Hpy99VI was deemed unlikely following the finding of no significant sequence variation in its coding sequence (Supplementary Fig. 10).

**Figure 4.**
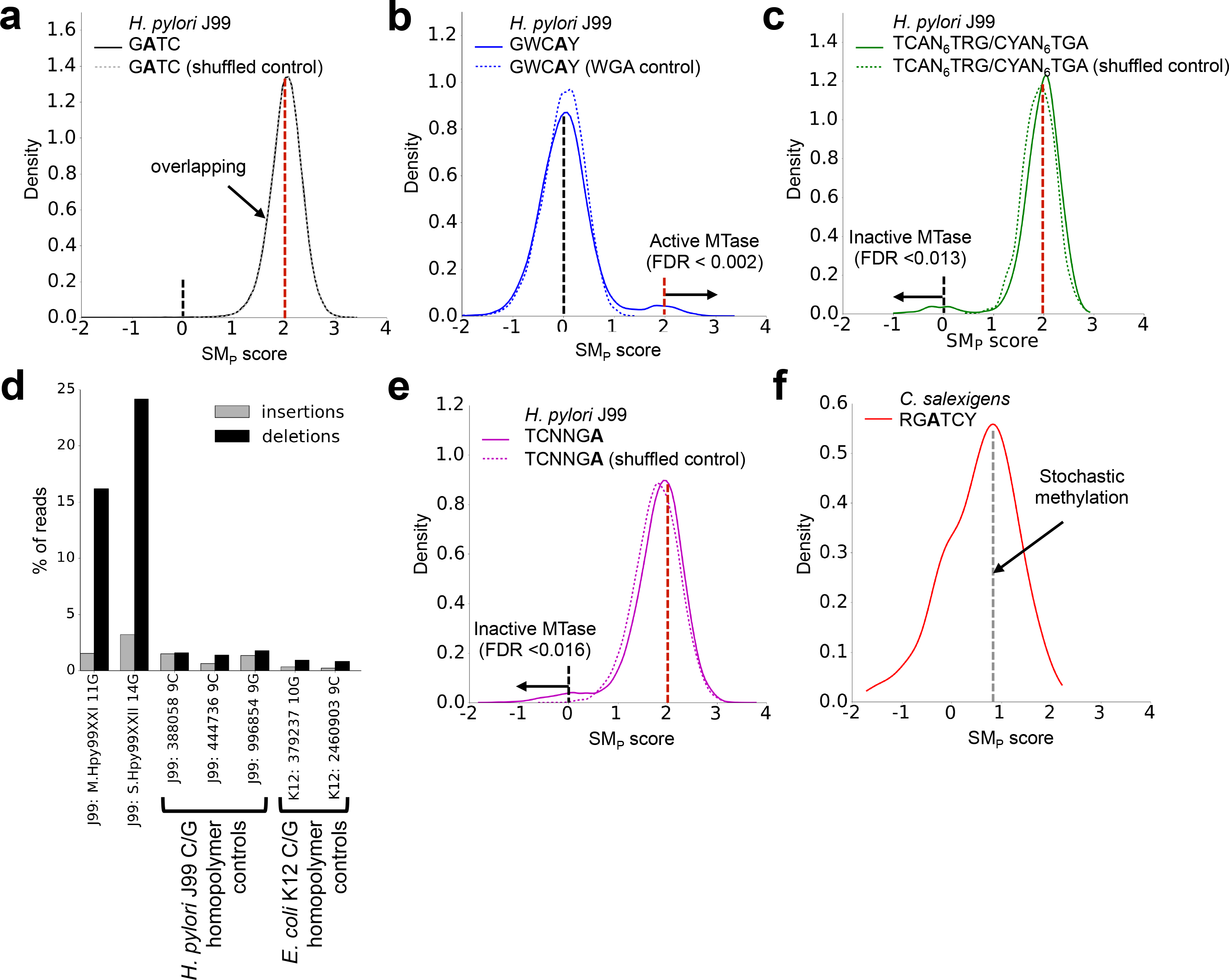
Single molecule, motif-pooled (SM_p_) approach for measuring MTase activity and detecting possible phasevarions. **(a)** SM_p_ distribution for *H. pylori* J99 motif GATC and its corresponding IPD-shuffled control. The identical unimodal distributions suggest a fully active MTase (as expected). **(b)** SM_p_ distribution for *H. pylori* J99 motif GWCAY and its corresponding WGA control SM_p_ distribution. The major peak around SM_p_ = 0 and small bump around SM_p_ ∼ 2 suggests that responsible MTase, M.Hpy99XXI, is mostly inactive, except for a small number that actively methylate GWCAY. Actively methylated molecules with SM_p_ scores > 2 have an FDR < 0.2%. **(c)** SM_p_ distribution of TCAN_6_TRG/CYAN_6_TGA (targeted by Hpy99XXII) in *H. pylori* J99, as well as its corresponding IPD-shuffled control SM_p_ distribution. SM_p_ analysis implicated this motif as having a potential role similar to that of GWCAY (i.e. small bimodality). In contrast to GWCAY, however, this SM_p_ distribution suggests the normally active MTase is sporadically inactivated in some cells. Non-methylated molecules with SM_p_ scores < 0 have an FDR < 1.3%. **(d)** High-accuracy sequencing with Illumina MiSeq and read-level analysis of insertion/deletion calls revealed the presence of significant variation in the lengths of two specific homopolymers in the coding sequences of M.Hpy99XXI and Hpy99XXII. The high percentage of deletions in these two genes stands apart from the deletion rates found in other C/G homopolymers from *H. pylori* J99 and *E. coli* K12, suggesting that this is not simply due to a higher rate of sequencing errors in homopolymer regions. **(e)** SM_p_ distribution of TCNNGA (targeted by M.Hpy99XVIII) in *H. pylori* J99, as well as its corresponding IPD-shuffled control SM_p_ distribution. This distribution of SM_p_ scores suggests a MTases behavior similar to that of TCAN_6_TRG/CYAN_6_TGA. Non-methylated molecules with SM_p_ scores < 0 have an FDR < 1.6%. (f) SM_p_ distribution for the *C. salexigens* motif RGATCY. The major peak near SM_p_ ∼ 0.9 indicates that the IPDs sampled for each molecule reflect a mixture of both non-methylated (IPD≈0) and methylated (IPD≈2) motif sites, suggesting stochastic methylation as the primary source of epigenetic heterogeneity for this motif.

Next, we applied the same analysis to a motif, GWCAY, targeted by an MTase that is known to be phase-variable in *H. pylori* J99. The full-length transcription of the gene encoding the modification (M) subunit of this type-III R-M system Hpy99XXI depends on the length of a specific 10-12G homopolymer^23^. Modification of the GWCAY motif is normally undetectable in *H. pylori* J99^23^, suggesting that the status of the 10-12G homopolymer in the isolate results in mainly inactive M.Hpy99XXI proteins. However, as the SM_p_ score distribution reveals (Fig. 4b), there is a subpopulation of molecules with SM_p_≈2, indicating the presence of a small fraction of cells containing the active form of M.Hpy99XXI in the *H. pylori* J99 culture. MiSeq-based sequencing of the same DNA supported this mechanism, revealing significant variation in the length of the *M.Hpy99XXI* homopolymer with respect to our reference assembly (Supplementary Fig. 11).

Hpy99XXII is another R-M system in *H. pylori* J99 that is known to contain phase-variable components. Instead of being found in the M subunit, the homopolymer-driven phase-variation in this system is found in the specificity (S) subunit, which targets the TCAN_6_TRG/CYAN_6_TGA motif. The presence of a 14G homopolymer in *S.Hpy99XXII* has been shown to result in a full-length specificity protein and active methylation of TCAN_6_TRG/CYAN_6_TGA^23^. Although our reference assembly contains the 14G homopolymer, we observe a small subpopulation of molecules with SM_p_≈0 (Fig. 4c), indicating that there are cells within the *H. pylori* J99 culture in which methylation by the Hpy99XXII system is not occurring. MiSeq sequencing confirmed the presence of significant length variation in this *S.Hpy99XXII* homopolymer in the culture (Supplementary Fig. 12).

The accuracy of sequencing homopolymeric regions, especially C/G stretches, is reduced with most sequencing platforms. To ensure that the observed insertions and deletions were not simply due to sequencing errors, we searched for length variation in other C/G homopolymers from both the same *H. pylori* J99 and *E. coli* K12 genomes that are not known to undergo phase variation (Fig. 4d). The number of read-level deletions observed in the *M.Hpy99XXI* (GWCAY) and *S.Hpy99XXII* (TCAN_6_TRG/CYAN_6_TGA) genes are markedly higher (>15%) than those observed in the control C/G homopolymers (<3%) observed in the control homopolymers (9- and 10-mers, the longest in the same *H. pylori* strain and the *E. coli* K12 genome), which is unlikely just due to the difference in homopolymer lengths. This suggests the homopolymer length variation, rather than sequencing error, is driving the variation present in these two methylation motifs. Interestingly, although a similar distribution of SM_p_ scores was observed for both the TCAN_6_TRG/CYAN_6_TGA (Fig. 4c) and TCNNGA (Fig. 4e) motifs, no sequence variation was observed in or around the coding region of the genes involved in the Hpy99XVIII R-M system responsible for the methylation of TCNNGA (Supplementary Fig. 13), suggesting that regulatory elements outside the coding region (genetic or epigenetic) may be responsible for the observed fraction of non-methylated molecules.

**Stochastic methylation in C. salexigens**. In contrast to GWCAY, TCAN_6_TRG/CYAN_6_TGA, and TCNNGA in *H. pylori* J99, we found no evidence from the SM_p_ scores supporting the existence of phase-variable MTase activity with the RGATCY motif of *C. salexigens* (Fig. 4f). Rather than showing a component of the SM_p_ score distribution centered on SM_p_≈2, we see a primary component centered on SM_p_≈0.9, indicating that the motif-pooled IPD values used to calculate each molecule’s SM_p_ score originated from a mixture of both non-methylated (IPD≈0) and methylated (IPD≈2) sites. This observation, combined with a lack of detected sequence variation in the coding sequence of the RGATCY-targeting M.CsaI gene (Supplementary Fig. 14) and the fact that M.CsaI is an orphan methyltransferase without corresponding RE^36^, indicates that stochastic methylation by M.CsaI is the likely mechanism driving the significant SM_SN_ heterogeneity observed in Figure 3b. Therefore, it is plausible that M.CsaI in *C. salexigens* has a stochastic methylation activity (∼60%) in each single cell, another type of global epigenetic heterogeneity in a bacterial population that differs from phase variation.

**Other motifs**. We also applied the long library-based SM_p_ analysis to the other motifs in the *H. pylori* J99 and *C. salexigens* methylomes, but did not observe SM_p_ score distributions suggesting additional phase-variable MTases (Supplementary Fig. 15). Generally, the power of long library-based SM_p_ analysis increases with the number of unique motif sites in the sequencing molecule, as shown in an ROC summary (Supplementary Fig. 16). 4-mer and 5-mer motif sites are often present at sufficient density in genomes such that the current read length of SMRT sequencing (mean 8,500bp, 95% percentile 20,000bp) can cover enough distinct sites to accurately assess the active/inactive state of the MTase Some long motifs have much higher frequencies than expected, such as the TCAN_6_TRG/CYAN_6_TGA motif in *H. pylori* J99, which has an average frequency of 10.8 per 10 kilobases, over two-fold higher than the frequency expected by chance for a k-mer of equal size (Supplementary Fig. 17). Therefore, we suggest using the actual number of 6mA sites in each molecule as a threshold for SM_p_ analysis instead of basing the threshold on motif length alone.

## Discussion

Here, we present the first systematic framework for single molecule-level characterization of epigenetic heterogeneity in bacterial methylomes using SMRT sequencing. We demonstrate the enhanced resolving power of our approach in analyses of seven methylomes that show distinct types of epigenetic heterogeneity in bacterial populations. The short library-based SM_SN_ method enables accurate estimation of the fractions of methylated and non-methylated motif sites without the use of subjective or *ad hoc* thresholds, even at low levels of genomic sequence coverage, such as sample-limited *in vivo* isolates. Furthermore, the robust separation between methylated and non-methylated sites based on SMsn scores at low coverage also suggests the possibility of *de novo* discovery of methylation motifs from SMRT sequencing data at a coverage that is much lower than the current standard. The second SM_p_ method leverages long library sequencing and single molecule motif pooling in order to phase methylation states and discriminate between different mechanisms that drive epigenetic heterogeneity in a bacterial population. By surveying phased MTase activity at a single molecule-level, the SM_p_ scores are able to reveal small fractions of cells that contain active and inactive MTases in population.

Together, the high resolution for surveying MTase activity provided by these complementary methods is not achievable using the existing, population-based analysis approaches. We therefore expect their application to improve the analysis and interpretation of bacterial methylomes. Furthermore, integration of this framework with other single molecule- or single cell-level data, such as RNA and protein expression, will enable a more detailed understanding of the functions of DNA methylations in bacterial physiology.

SMRT sequencing provides a unique opportunity to *de novo* detect more than twenty different types of DNA modifications^32,46,51^, including all the three major types of methylations in the bacterial kingdom, but also poses special challenges that previous second-generation sequencing techniques have not been forced to address. Specifically, for most second-generation sequencing techniques, the fidelity of single molecule-level analysis is often taken for granted because of the high sequencing accuracy during a single pass over the template. However, for SMRT sequencing, the accuracy of base calling and detection of DNA modifications fundamentally depends on the number of repeated observations for each single molecule (circular consensus). Therefore, given a fixed average read length in SMRT sequencing, there is a tradeoff between library size and accuracy in both base calling and methylation detection. The methods proposed here provide an example of how to effectively leverage the unique features of SMRT sequencing using a combination of library designs. Shorter libraries have higher accuracy in detecting methylation at the level of single molecules and individual nucleotides, but are less informative when trying to determine the activity states of MTases along the full length of a single read. The short library-based method provides a highly accurate estimation of the global fraction of methylated and non-methylated sites, as well as detection of non-methylated sites at single molecule resolution. Alternatively, longer libraries enable the assessment methylation states along long templates, thereby facilitating inferences regarding MTase activity states, but do not provide reliable single molecule, single nucleotide-level methylation calls. The hybrid approach, using the two methods, can yield unprecedented insight into bacterial methylomes. Furthermore, the general methodology used in the design of these two methods can be used as a template to design analytic approaches for forthcoming third-generation real-time sequencing techniques^41,52^.

Collectively, the methods presented here will facilitate new studies and insights into the heterogeneity and significance of methylation within bacterial populations and mechanisms that underlie it. Finally, although the current study focused on cultures of single bacterial strains, the single molecule resolution methods proposed here can also be applied to mixed populations of bacteria. Such samples could include diverse clinical isolates, including samples that contain a pathogen in low abundance or even diverse microbiome samples. The methods are also applicable outside the bacterial kingdom, such as in the analysis of DNA from human mitochondria or viruses, both of which present significant genetic and epigenetic heterogeneity.

## Acknowledgement

We thank Jonas Korlach and Richard Roberts for providing access to raw sequencing data as well as secondary analysis of samples from Murray et al. We also thank Lucy Shapiro and Harley McAdams for providing access to raw sequencing data of samples from Kozdon et al. The work was supported in part by R01-GM63270 from the National Institutes of Health, the Diane Belfer Program for Human Microbial Ecology, and the Daniel Ziff Fund. This work was also supported in part through the computational resources and staff expertise provided by the Department of Scientific Computing at the Icahn School of Medicine at Mount Sinai.

## Online Methods

### Bacterial strains and culture conditions

The *Escherichia coli* C227 sample was isolated from a 64-year-old woman from Hamburg, Germany, who was hospitalized in Copenhagen, Denmark after presenting with bloody diarrhea. A full description of the DNA extraction and sequencing procedures is provided in Rasko *et al*^1^.

*Helicobacter pylori* J99 was originally isolated by Blaser lab from an American patient with duodenal ulcer^2^ and sequenced in 1999^3^. *H. pylori* cells (50 μl) of J99 frozen stock from Blaser lab were first spotted on trypticase soy agar (TSA) plates with 5% sheep blood (TSA, BBL Microbiology Systems, Cockeysville, MD) at 37°C with 5% CO2 for 48-hour incubation, and then spread on new TSA plates in the same conditions^4^. After 24-h incubation, *H. pylori* cells were collected in 1.0 mL phosphate-buffered saline (PBS, pH 7.4), and centrifuged at 800 g for 5 min. Bacterial genomic DNA was prepared using the Wizard Genomic DNA purification kit as following manufacturer’s instructions (Promega, Madison WI.). DNA concentration was measured by Nanadrop 1000 spectrophotometer (Thermo Scientific, Rockford, IL).

DNA for *Chromohalobacter salexigens* strain 1H11 is ordered from DSM (http://www.dsmz.de/catalogues/details/culture/DSM-3043.html).

The procedures for culturing, synchronizing, and isolating genomic DNA from *Caulobacter crescentus* are described by Kozdon *et al*^5^. The DNA isolation procedures for *Geobacter metallireducens, Campylobacter jejuni* 81-176, & *C. jejuni* NCTC 11168 are described in Murray *et al*^6^.

### Whole genome amplification (WGA)

The Qiagen REPLI-g amplification kit was used to perform whole-genome amplification to exclude epigenetically modified bases. The method produced micrograms of DNA from 50 ng of input genomic DNA, following the manufacturer’s guidelines and 10 hours of amplification time at 30^o^C followed by deactivation at 65^o^C for 3 minutes.

### SMRT sequencing

Long (20,000bp) insert DNA library preparation and sequencing was performed according to the manufacturer’s instructions and reflects the P5-C3 sequencing enzyme and chemistry, respectively. Upon completion of library construction, samples were validated as ∼20,000bp using an Agilent DNA 12000 gel chip. All isolate libraries were sufficient for additional size selection to remove any SMRTbells < 7,000 bp. This step was conducted using Sage Science Blue Pippin 0.75% agarose cassettes to select library in the range of 7,000 bp −50,000 bp. This selection is necessary to narrow the library distribution and maximize the SMRTbell sub-readlength. 11–23% of the input libraries eluted from the agarose cassette and was available for sequencing. For all cases, this yield was sufficient to proceed to primer annealing and DNA sequencing on the PacBio RSII machine. Primer was then annealed to the size-selected SMRTbell with the full-length libraries (80°C for 2 minute 30 followed by decreasing the temperature by 0.1°/s to 25°C). The polymerase-template complex was then bound to the P5 enzyme using a ratio of 10:1 polymerase to SMRTbell at 0.5 nM for 4 hours at 30°C and then held at 4°C until ready for magbead loading, prior to sequencing. The magnetic bead-loading step was conducted at 4°C for 60-minutes per manufacturer’s guidelines. The magbead-loaded, polymerase-bound, SMRTbell libraries were placed onto the RSII machine at a sequencing concentration of 75 pM and configured for a 180-minute continuous sequencing run.

For all short (250bp) insert library preparations, similar methodology was used, except shearing was done using a Covaris microtube ultrasonication and all AMPure XP purification steps were done using a 1.8X volume ratio. Libraries were completed without the size selection step used for the long insert libraries. Similar procedures were followed for sequencing, except that diffusion-based loading was used instead of magbead loading.

### Illumina MiSeq sequencing and analysis (*C. salexigens, H. pylori*)

Illumina-based MiSeq whole-genome libraries were prepared with an insert size of 600 bp as assessed by Agilent Bioanalysis using standard Illumina adapters and 8 PCR cycles. 2 x 350 bp paired-end sequencing was then conducted using version 3 commercial kits to assure the longest readlength possible.

The *E. coli* K12 MiSeq reads used for analysis of homopolymer indel rates (Figure 4d) was downloaded from http://www.illumina.com/systems/miseq/scientific_data.ilmn. The sequencing reads from the *E. coli* K12, *C. salexigens,* and *H. pylori* J99 MiSeq runs were aligned to their respective references using *bwa mem*^13^ with its default parameters. The resulting alignments were processed with the *samtools* package^14^ to obtain pileups for each genomic position. Read-level mismatch and insertion/deletion calls were obtained from the pileups by counting the number of occurrences of each type of variant at each genomic position.

### Bioinformatics analysis

The details of the bioinformatics analysis are described below. The methods were implemented in a stand-alone software package and will be released at Bitbucket upon publication of the manuscript, or upon request to the corresponding authors.

### Filtering subreads and preprocessing

An initial filtering step removes all subreads with ambiguous alignments (MapQV<240), low accuracy (<80%), or short aligned length (<100 bases). Next, because sequencing errors in the subreads are likely to introduce noise into the IPD distribution that are being used to infer methylation status, an additional filtering step removes from further analysis the subread IPD values from the positions +1:-1 on either side of any errors with respect to the reference sequence. This removes a substantial number of IPD values from consideration, but those that remain are minimally impacted by sequencing errors. Furthermore, the first ten and last fifteen bases from each subread are removed from analysis due to the potential for bias in the IPD values near the transition between template and adapter sequences. Finally, subread normalization was used to account for potential slowing of polymerase kinetics over the course of an entire read (consisting of many subreads) as was done in previous studies^7^.

### Population-level detection of methylation states (Agg_SN_)

In accordance with the method described by Flusberg *et al*^8^, the IPD values from all native molecules were aggregated according to their strand and mapped genomic position. Similarly, the IPD values from all the whole-genome amplified (WGA) molecules were aggregated according to their strand and mapped genomic position. The Agg_SN_ score per strand/position represents the log ratio of the native IPD mean to the WGA IPD mean.

### Single-molecule single-nucleotide detection of methylation states (SM_sn_)

Considering each native molecule separately, the IPD values for a given motif that remain after the initial filtering are grouped by their strand and mapped genomic position. Following natural log conversion of the set of IPD values for the molecule/strand/position, the mean value is calculated. Contrary to the treatment of native IPD values, the WGA IPD values are aggregated across all molecules covering each strand and mapped genomic position. This aggregation is done because all molecules are expected to be free of any DNA methylations due to the amplification process^7 8^’. The SM_sn_ is calculated by subtracting the WGA strand/position-matched mean IPD value from this native molecule/strand/position-specific mean IPD value. This is essentially a log scale IPD ratio^8^, which approximately follows a normal distribution^9^.

### Estimating sensitivity and specificity of SM_sn_ based detection of DNA methylations

To quantify the sensitivity and specificity for single molecule, single nucleotide-level detection, we tested this framework on methylated 5’-CTGC<u>A</u>G-3’ sites in a native *E. coli* 0104:H4 C227-11strain^7^ (C227), and a matching whole genome amplified (WGA) sample in which the methylation sites were erased. In the analysis, we assumed 100% of the CTGCAG sites were methylated in the native DNA to leverage the large number of motif sites in the native bacterial genome. This is expected to provide a more robust estimation of sensitivity and specificity compared to the use of short DNA oligos with the more limited diversity of expanded local sequence contexts flanking any given methylation motif site^9,10^. Generally speaking, given that CTGCAG sites are part of an active type-II RM system, it is reasonable to assume the vast majority of sites are methylated to prevent restriction of the host DNA. However, as shown in recent studies^5,7, 11^, a small number of motif sites (even those that are part of RM systems) are found to be non-methylated, likely due to competitive binding between the MTase and other DNA binding proteins or to the transient non-methylated window that is just prior to the replication fork. Therefore, the sensitivity estimated in the above analysis represents the lower bound of the actual sensitivity of our method.

### Approximate SM_sn_ scores for C. crescentus

WGA sequencing was unavailable for analysis in this study, so the approximate SM_sn_ score shown in Figure 3c is simply the native molecule/strand/position mean IPD value (without subtracting the strand/position-matched WGA mean IPD value). This approximate SM_sn_ score is susceptible to IPD biases introduced by local sequence contexts, but provides a sufficient ability to resolve the modified and non-modified components in the bimodal distributions as shown in Figure 3c.

### Single-molecule, motif-pooled detection (SM_p_)

First, considering each molecule separately, the native IPD values for a given motif that survive filtering are grouped by their strand only, irrespective of their mapped genomic positions. The natural log is taken for all values in this group and the mean calculated. Second, the strand/position-matched WGA IPD values are aggregated across all molecules as with the SM_sn_ method. However, in this case the WGA IPD values are additionally aggregated across each motif site covered by the native molecule in question. Then, these WGA strand- and motif site-matched IPD values are natural log-converted and their mean value is subtracted from the native molecule/strand-specific mean, resulting in the SM_p_ score.

### False discovery rate (FDR) estimation for identifying active/inactive molecules

To assess the significance of observed modified and non-modified fractions based on SM_p_ scores, we established negative control SM_p_ distributions in order to assess the FDR associated with a particular SM_p_ threshold. When assessing the significance of actively methylated molecule calls (e.g. for the GWCAY motif in *H. pylori* J99), the negative control SM_p_ distribution is generated by running the SM_p_ pipeline on WGA sequencing data. Alternatively, when assessing the significance of non-methylated molecule calls (e.g. for the TCAN_6_TRG/CYAN_6_TGA and TCNNGA motifs in *H. pylori* J99), the negative control SM_p_ distribution is created by randomly shuffling IPD values among molecules to disperse the non-modified IPD values for the motif of interest. Given that the number of non-modified IPD values is relatively low, this creates a control distribution of SM_p_ scores reflecting molecules that are mostly modified at the motif of interest.

### Estimation of modified fraction using Gaussian mixture modeling

The *mixture* module of the Python package *PyMix* was used to run the expectation maximization (EM) algorithm^12^ for Gaussian mixture estimation.

To evaluate the ability of EM with a Gaussian mixture model to estimate the modified fraction, we applied the algorithm to distributions of SM_sn_ scores that were generated from *in silico* mixing of WGA and native SMRT sequencing molecules. 100,000 total molecules sequenced from *E. coli* C227 were used for each specific mixture fraction, with the native fraction of molecules ranging from 5000 (5%) to 100,000 (100%), and the SM_sn_ score distributions for the motif CTGCAG was analyzed with the EM algorithm. To assess the stability of EM-mediated estimation of modified fraction at lower levels of genomic coverage (i.e. the number of total sequenced bases in relation to the genome size), we downsampled the SM_sn_ pipeline output for the *in silico* mixtures of WGA and native molecules. The EM algorithm was then applied to these downsampled mixed distributions.

The EM algorithm slightly, yet consistently, underestimates the native fraction in this mixture. In order determine whether any non-normality in the modified and non-modified SM_sn_ score distributions might be causing the EM algorithm to underestimate the size of the modified fraction, we created Gaussian mixtures consisting of two entirely simulated normal distributions. To represent the non-modified and modified SM_sn_ scores, the distributions were simulated with α=2 and α=0, respectively (σ=0.5 for each). For comparison, we also created mixtures using SM_sn_ values exclusively from WGA molecules where +2 was added to a subset of the SM_sn_ scores in order to get between 5% and 100% “modified” SM_sn_ values, the distributions of which will retain any non-normality found in the WGA SM_sn_ distribution. The EM estimations of modified fraction for these two types of simulated mixtures were both very similar and very accurate (Supplementary Figure 2), indicating that non-normality of SM_sn_ score distributions is not the reason for the observed slight underestimation of native fraction. Instead, this evidence suggests that stably non-modified CTGCAG sites in *E. coli* C227 are the reason for this phenomenon.

## Supplemental Figures

**Supplementary Figure 1:**
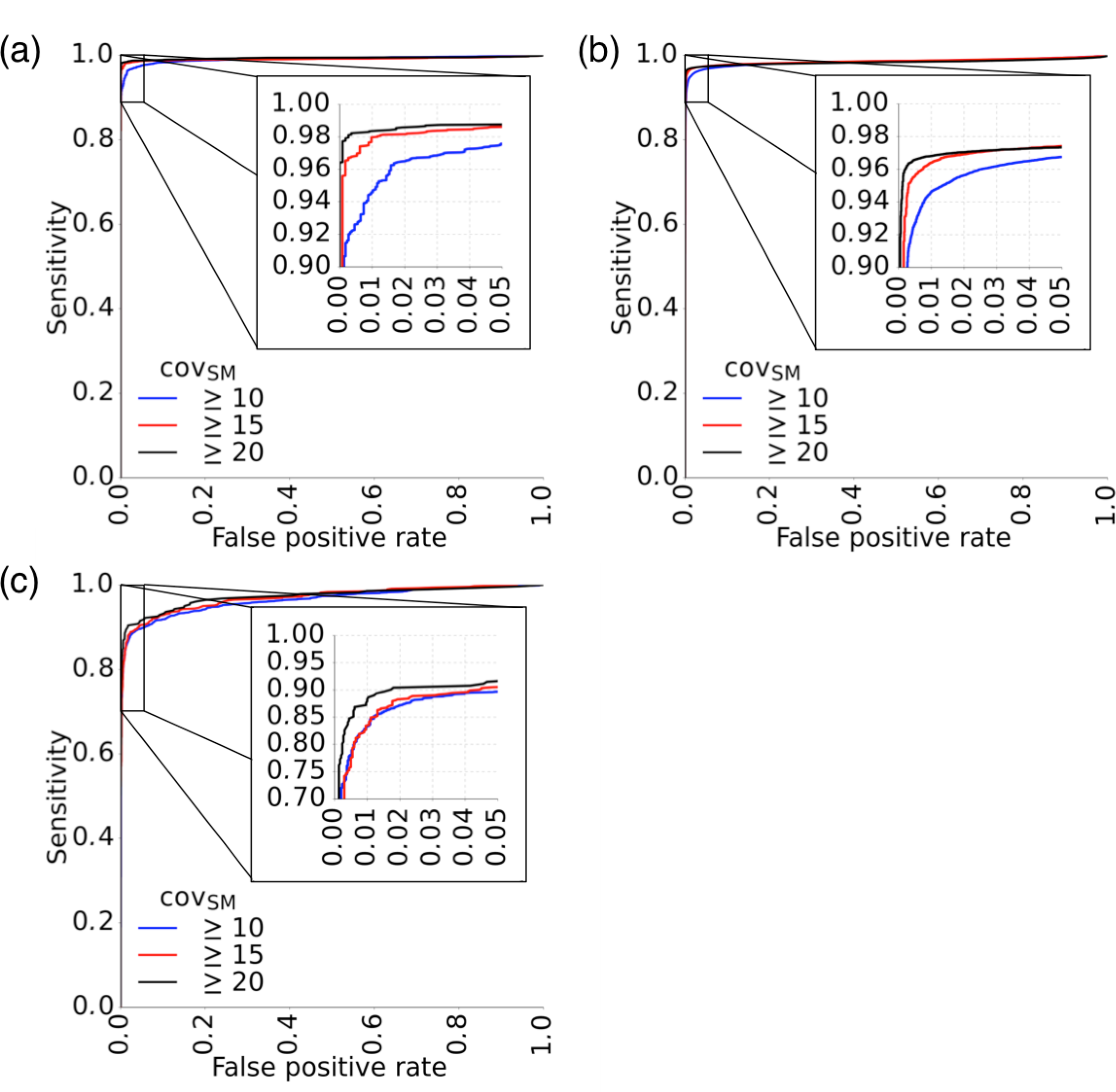
Sensitivity and specificity of the SM_sn_ method for detecting DNA modifications using three thresholds for minimum single-molecule coverage (cov_sm_): **(a)** 6mA modifications in the 5’-ACCACC-3’ motif in *E. coli* C227, **(b)** 6mA modifications in the 5’-GATC-3’ motif in *E. coli* C227, and **(c)** 4mC modifications in the 5’-CGWCG-3’ motif in *H. pylori* J99.

**Supplementary Figure 2:**
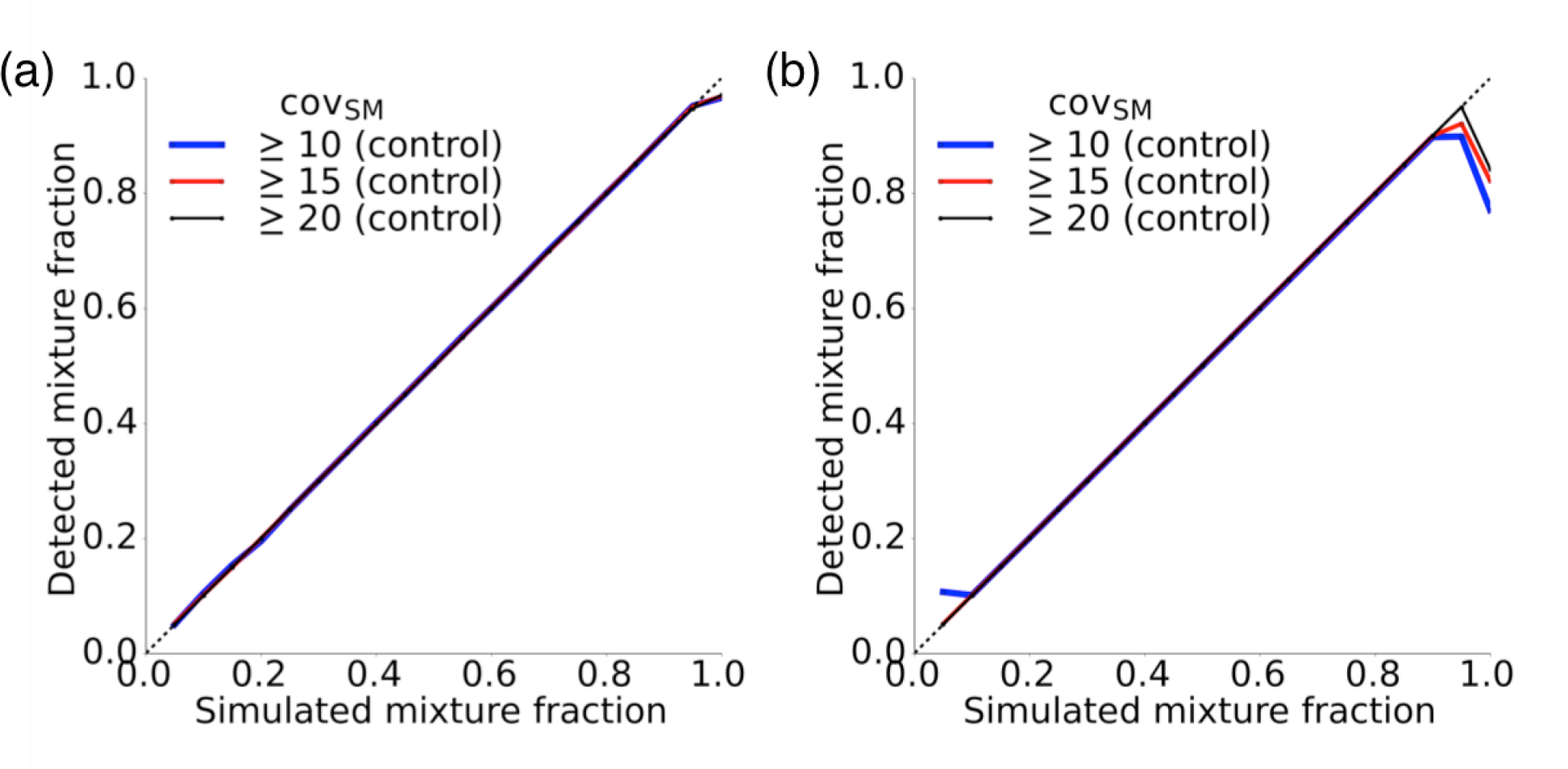
Two alternative approaches for generating simulated modified and non-modified distributions used to test the ability of the SM_Sn_ scores to estimate the size of the modified fraction. Instead of mixing varying proportions of WGA and native molecules, we instead **(a)** simulated two normal distributions centered around the mean SM_Sn_ scores for modified (SM_Sn_=0) and non-modified (SM_Sn_=2) adenine residues and varied their relative proportions, and **(b)** exclusively used WGA molecules, but simulated the presence of a modification by adding 2 to the WGA SM_Sn_ scores in a varying number of WGA molecules.

**Supplementary Figure 3:**
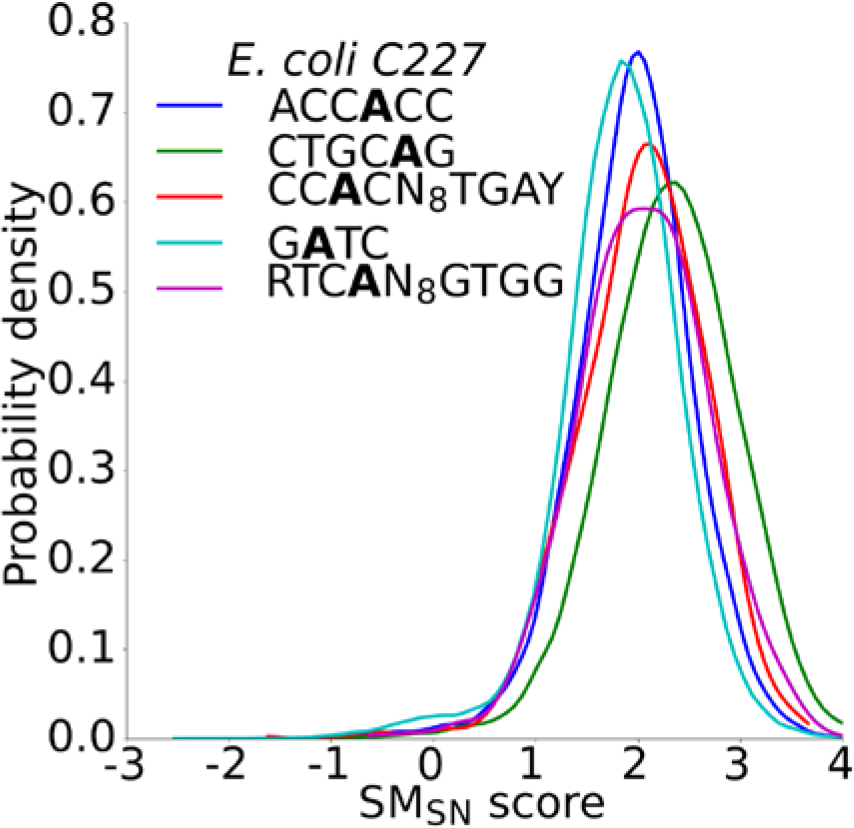
SM_sn_ score distributions (cov_sm_≥10) for all five 6mA motifs in *E. coli* C227.

**Supplementary Figure 4:**
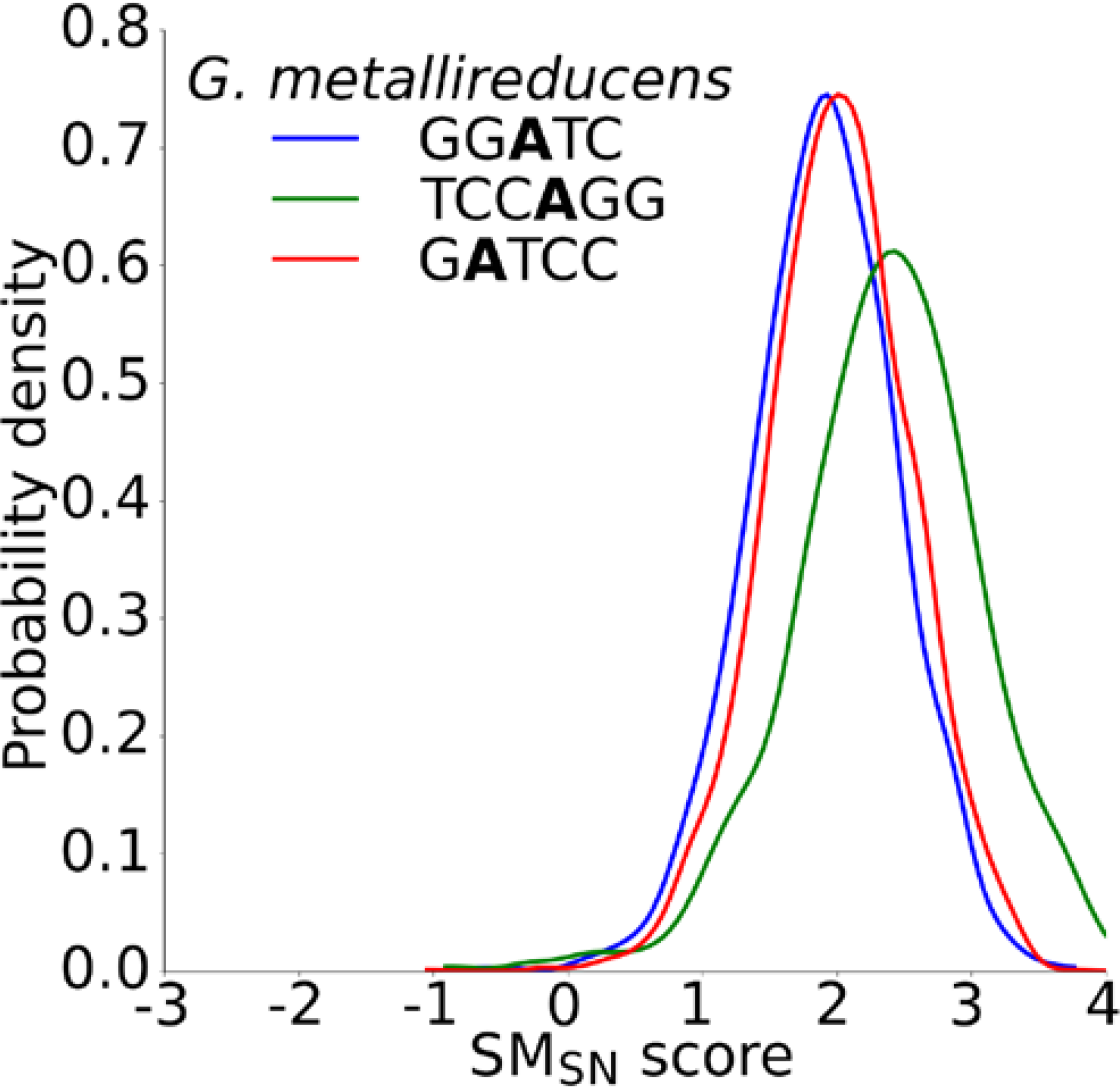
SM_sn_ score distributions (cov_sm_≥10) for all three 6mA motifs in *G. metallireducens.*

**Supplementary Figure 5:**
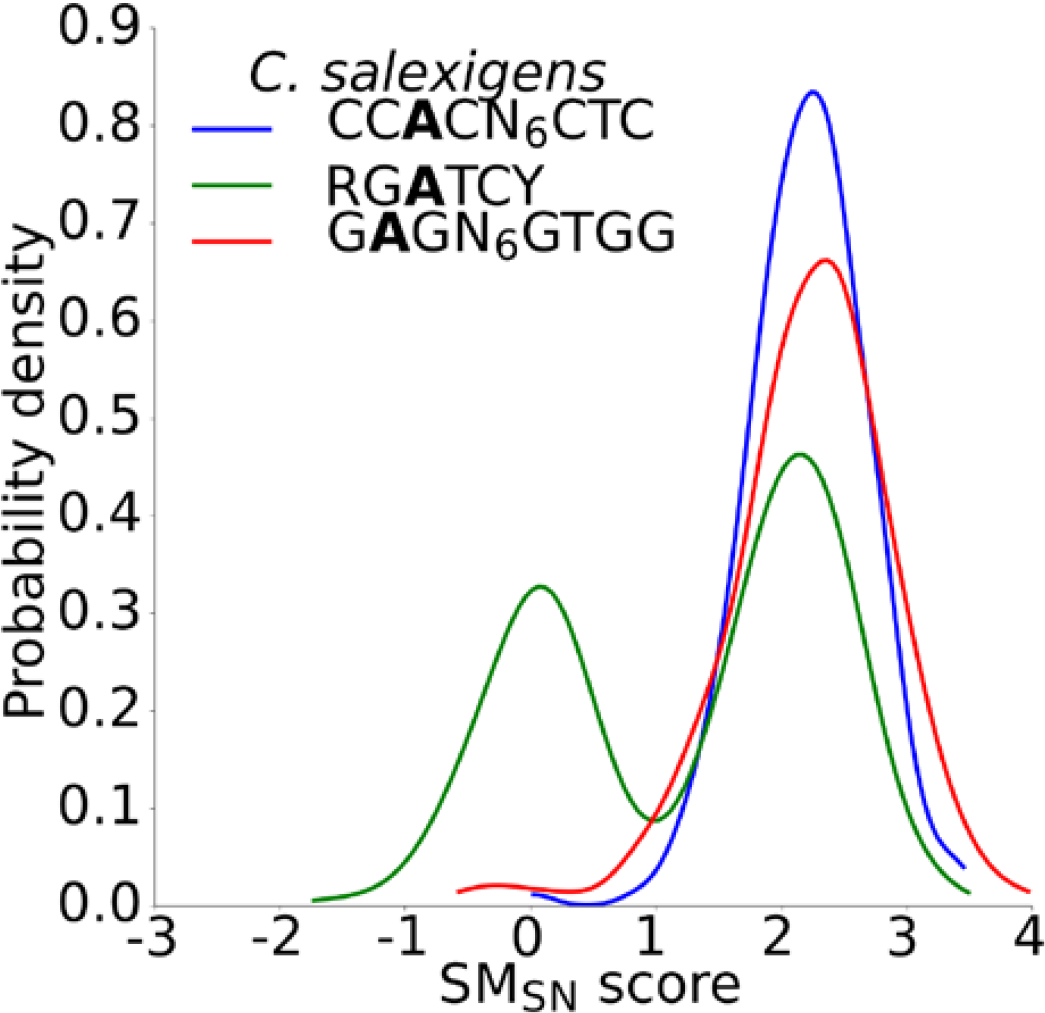
SM_sn_ score distributions (cov_sm_≥10) for all three 6mA motifs in *C. salexigens.*

**Supplementary Figure 6:**
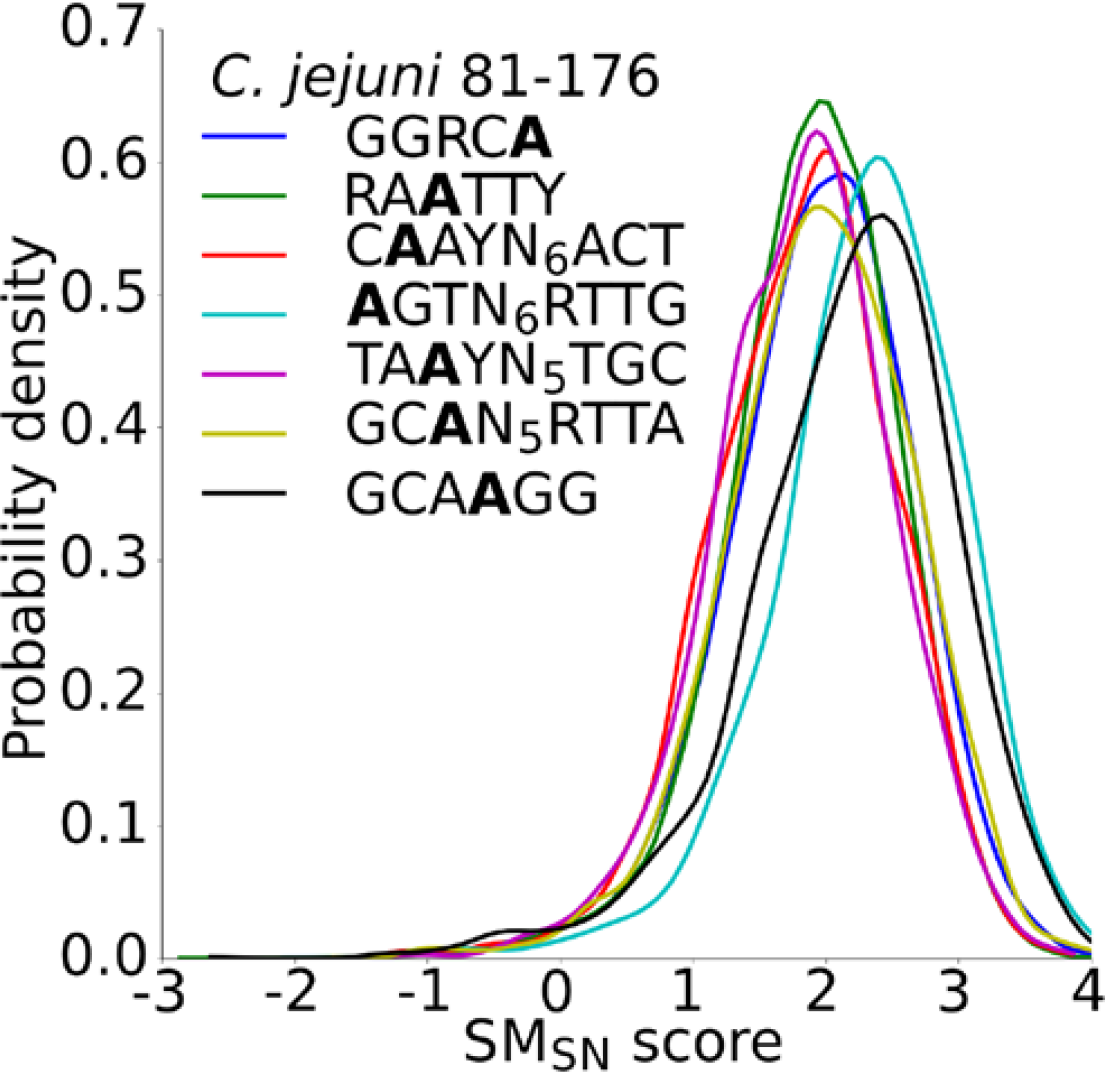
SM_sn_ score distributions (cov_sm_≥5) for all seven 6mA motifs in *C. jejuni* 81-176.

**Supplementary Figure 7:**
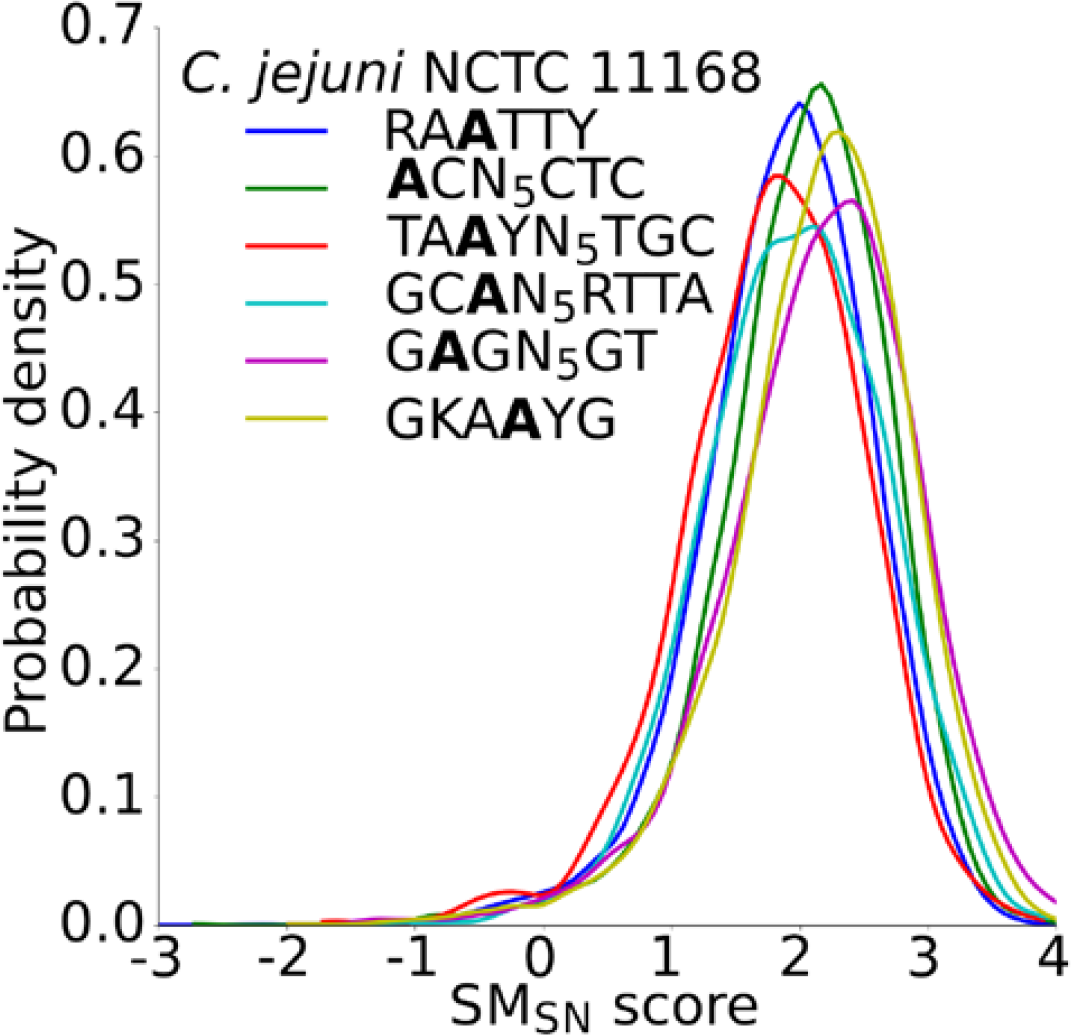
SM_sn_ score distributions (cov_sm_≥5) for all six 6mA motifs in *C. jejuni* NCTC 11168.

**Supplementary Figure 8:**
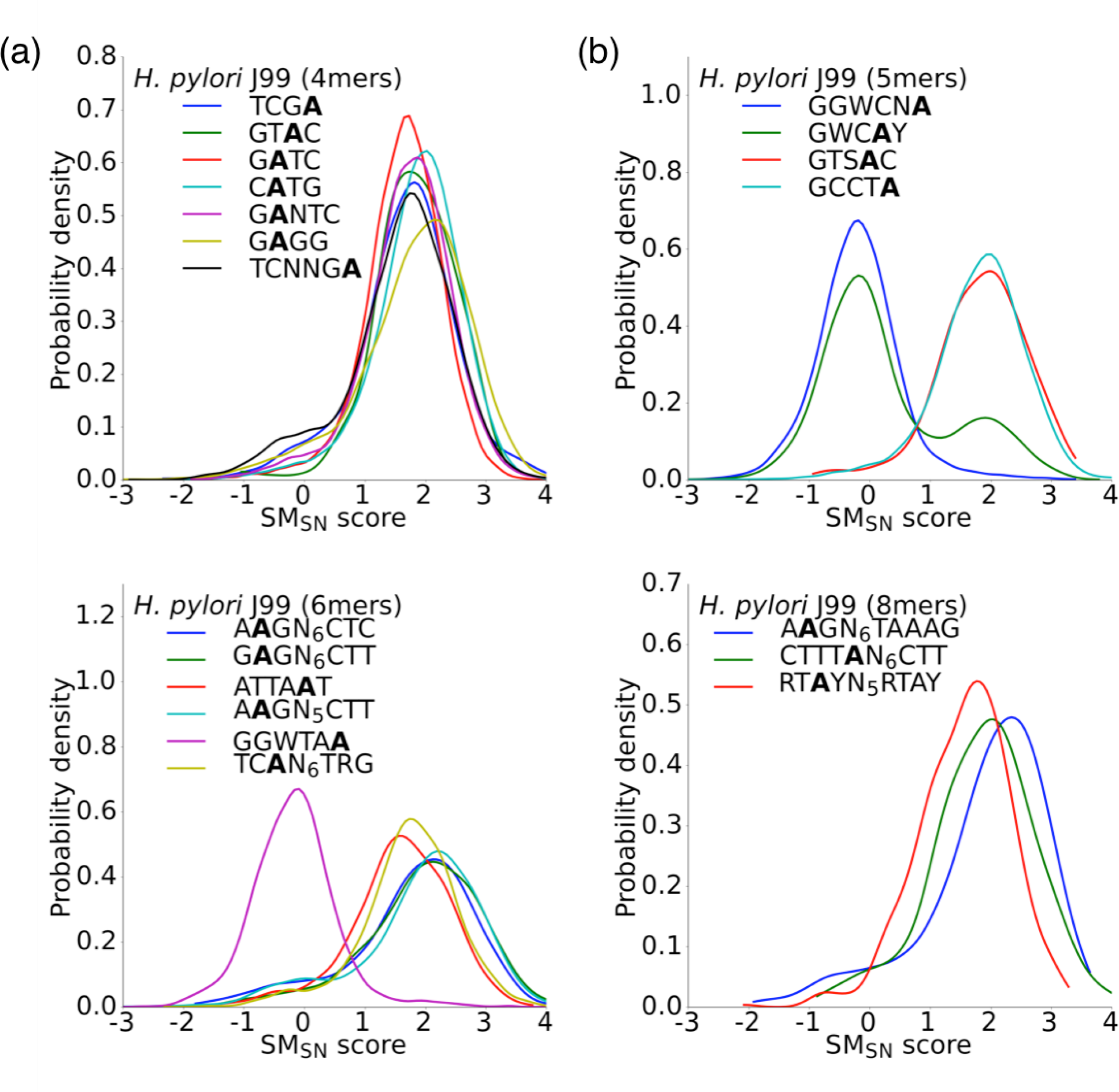
SM_sn_ score distributions (cov_sm_≥5) in *H. pylori* J99 for **(a)** all 4mer 6mA motifs, **(b)** all 5mer 6mA motifs, **(c)** all 6mer 6mA motifs, and **(d)** all 8mer 6mA motifs.

**Supplementary Figure 9:**
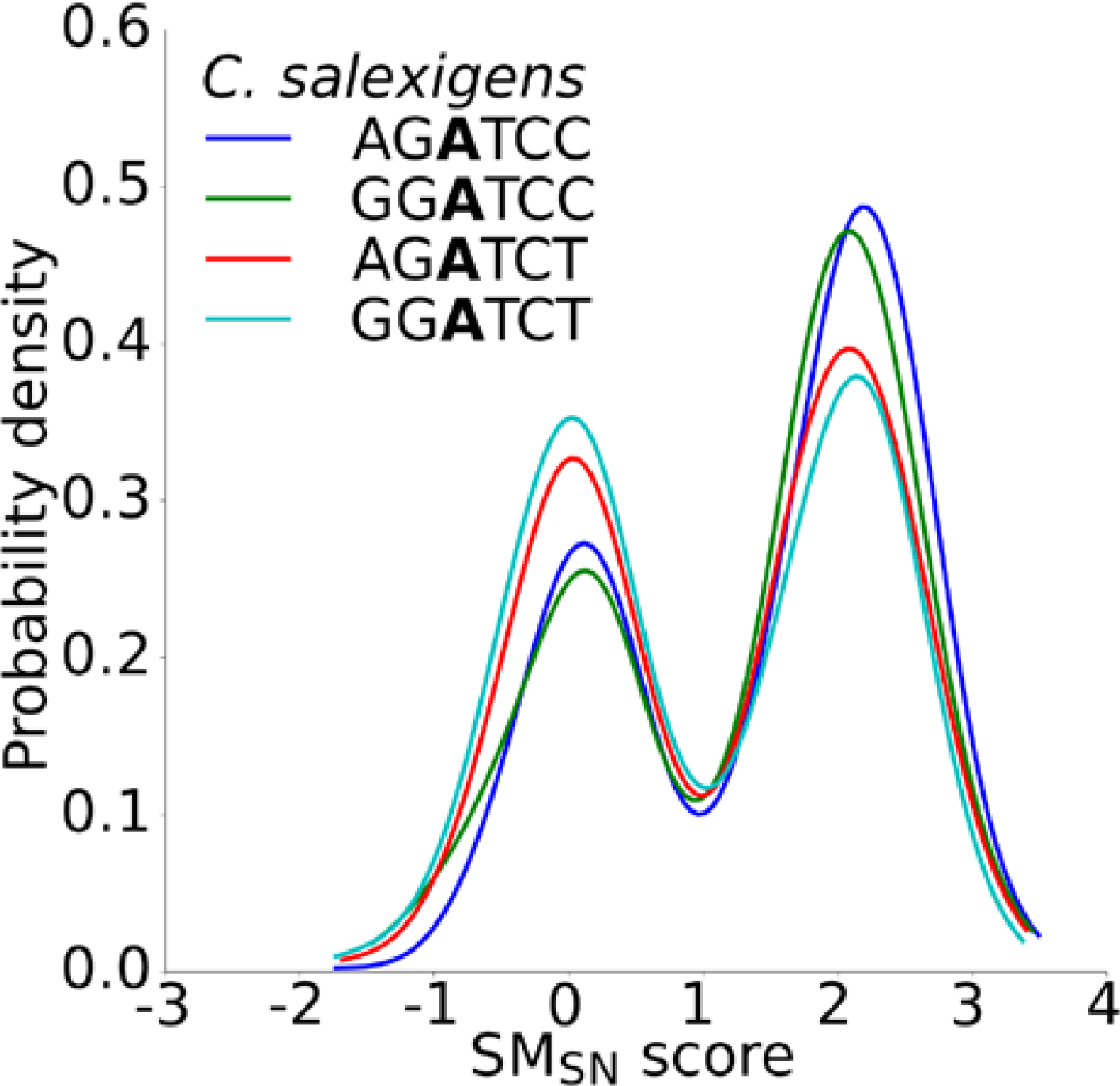
The SM_sn_ score distributions (cov_sm_ ≥10) for each specification of the degenerate 5’-RGATCY-3’ 6mA motif show similar levels of global heterogeneity.

**Supplementary Figure 10:**
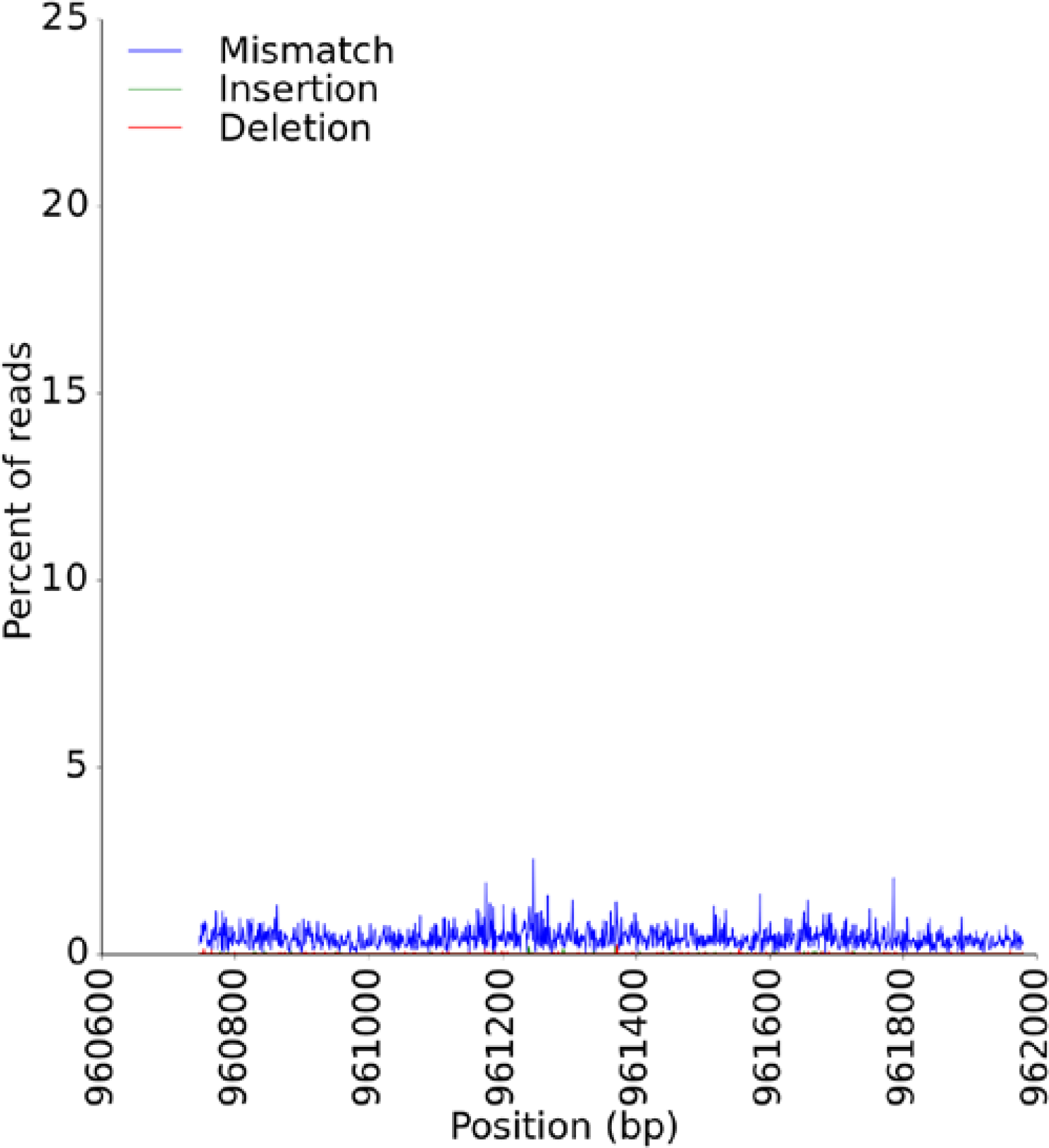
Read-level MiSeq mismatch and insertion/deletion calls in the coding region (+/-200bp) of the GATC-targeting *M.Hpy99VI* gene in *H. pylori* J99.

**Supplementary Figure 11:**
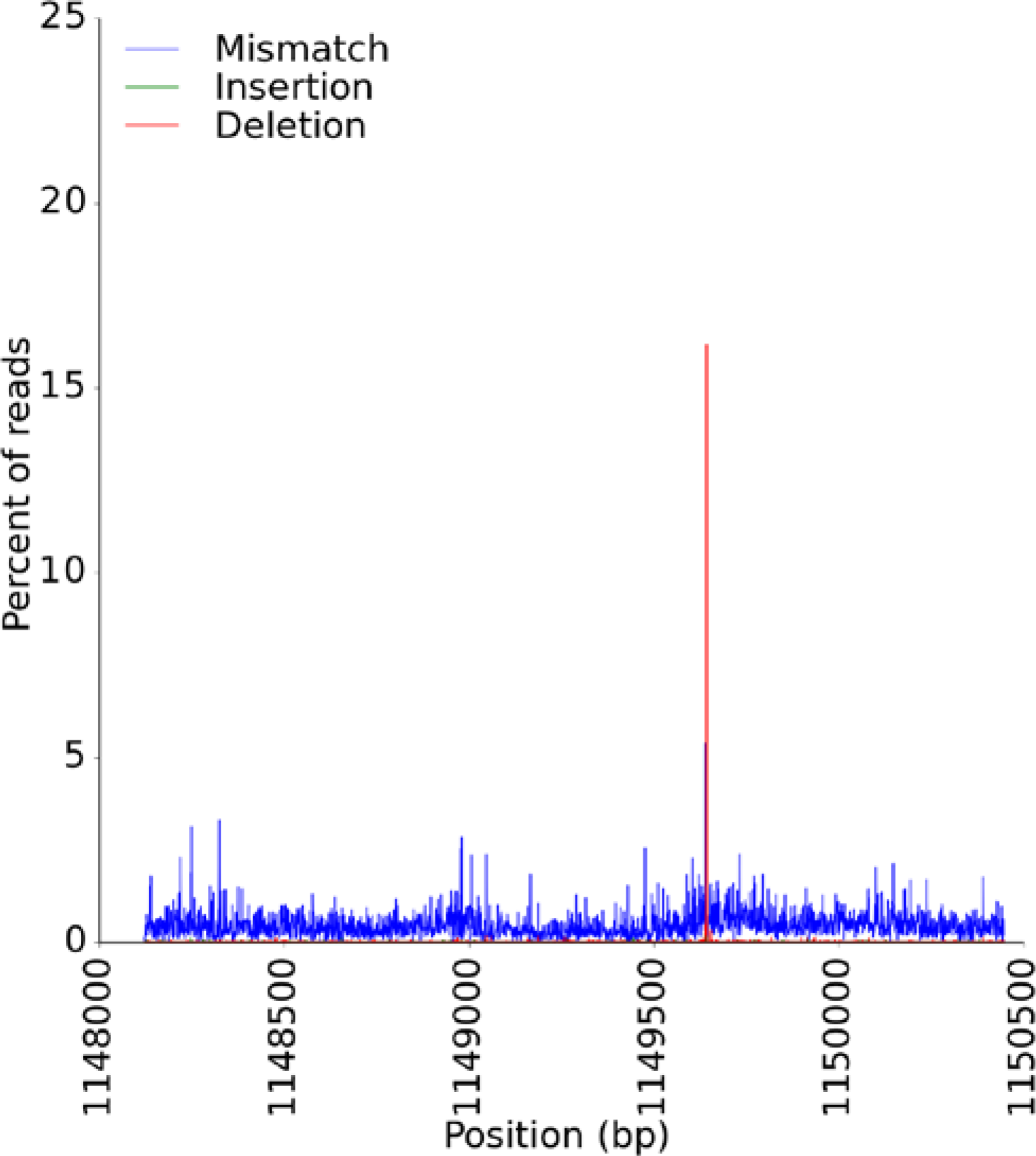
Read-level MiSeq mismatch and insertion/deletion calls in the coding region (+/-200bp) of the GWCAY-targeting *M.Hpy99XXI* gene in *H. pylori* J99.

**Supplementary Figure 12:**
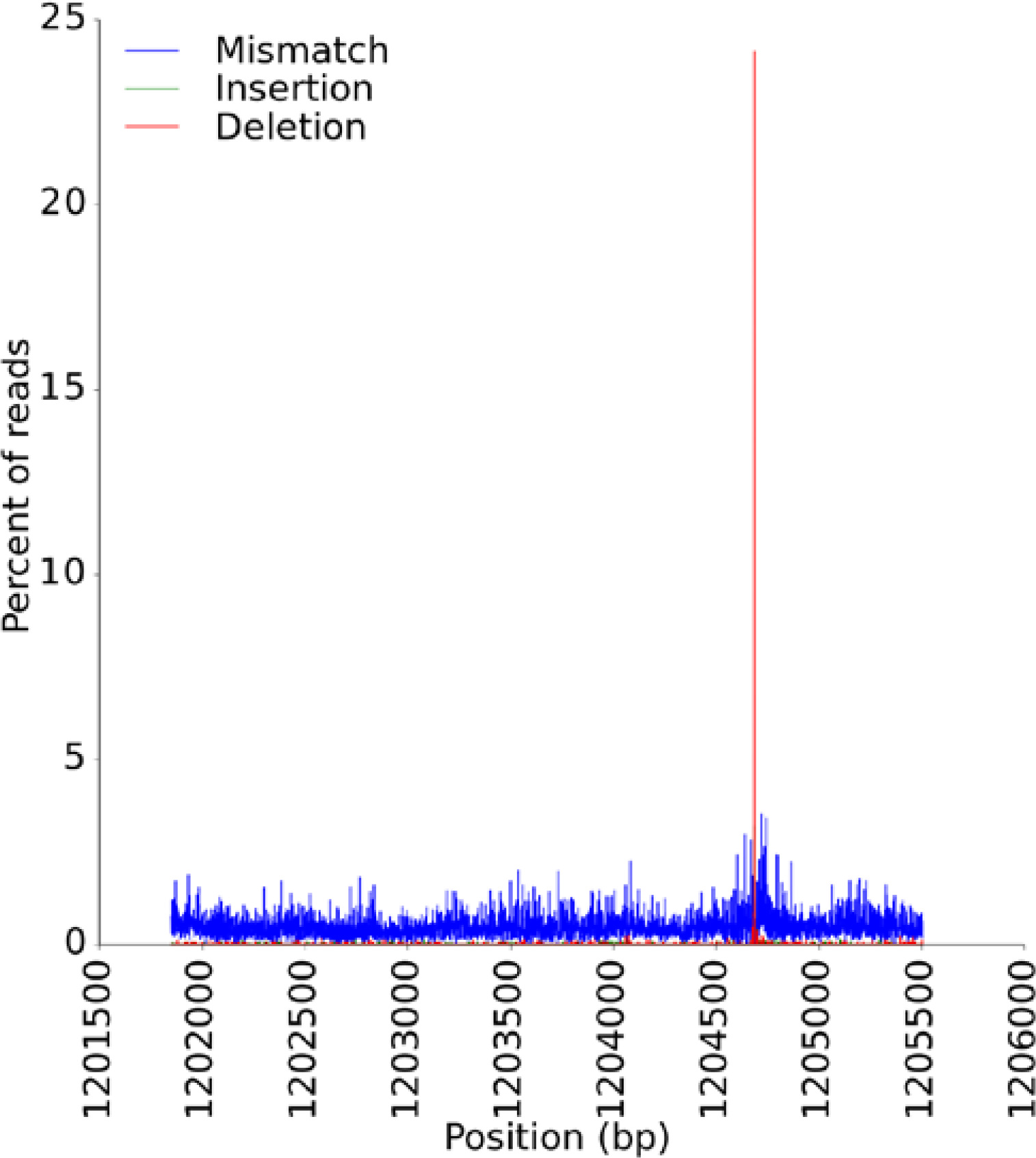
Read-level MiSeq mismatch and insertion/deletion calls in the coding region (+/-200bp) of the TCAN_6_TRG-targeting *Hpy99XXII* and *S.Hpy99XXII* genes in *H. pylori* J99.

**Supplementary Figure 13:**
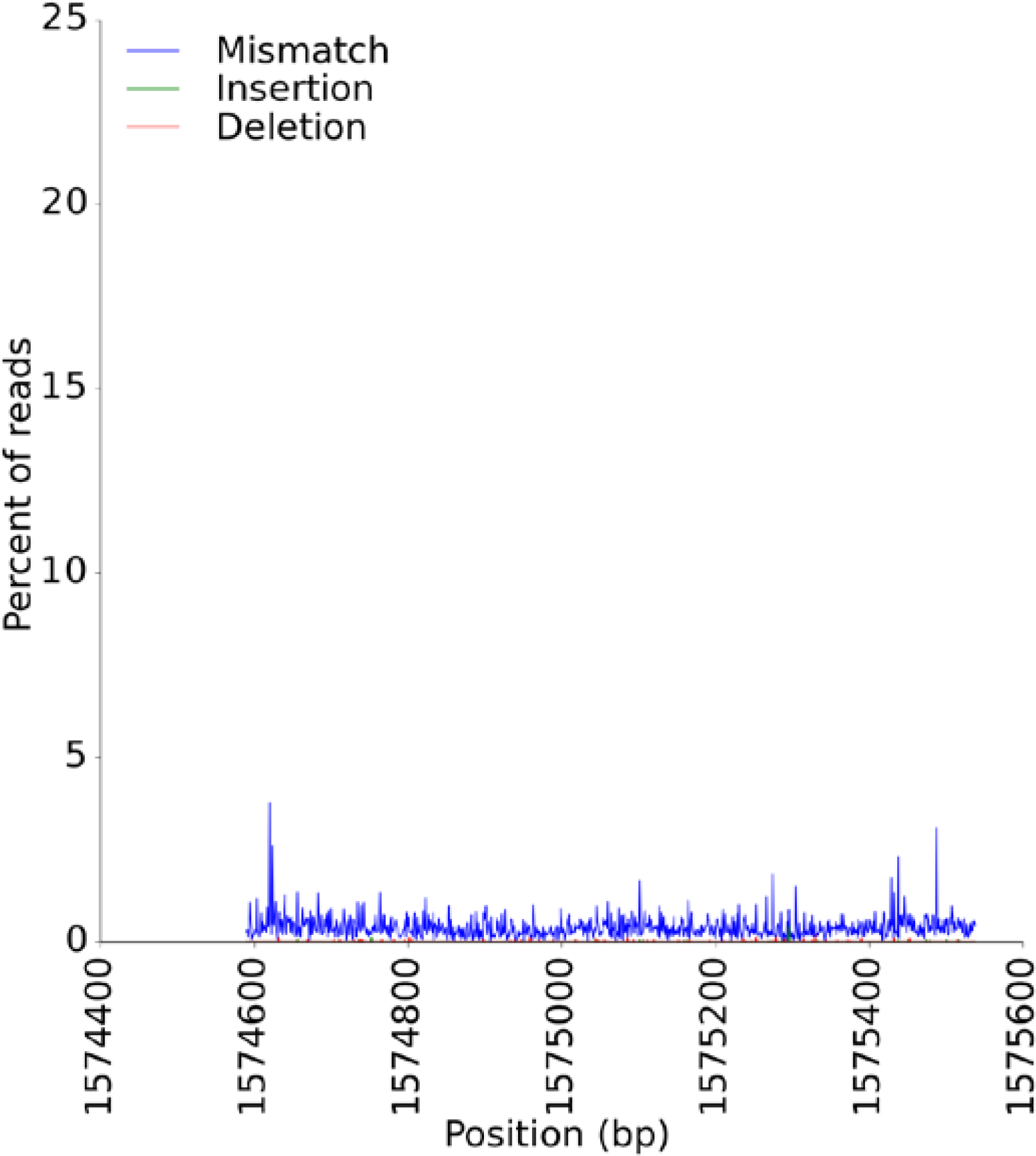
Read-level MiSeq mismatch and insertion/deletion calls in the coding region (+/-200bp) of the TCNNGA-targeting *M.Hpy99XVIII* gene in *H. pylori* J99.

**Supplementary Figure 14:**
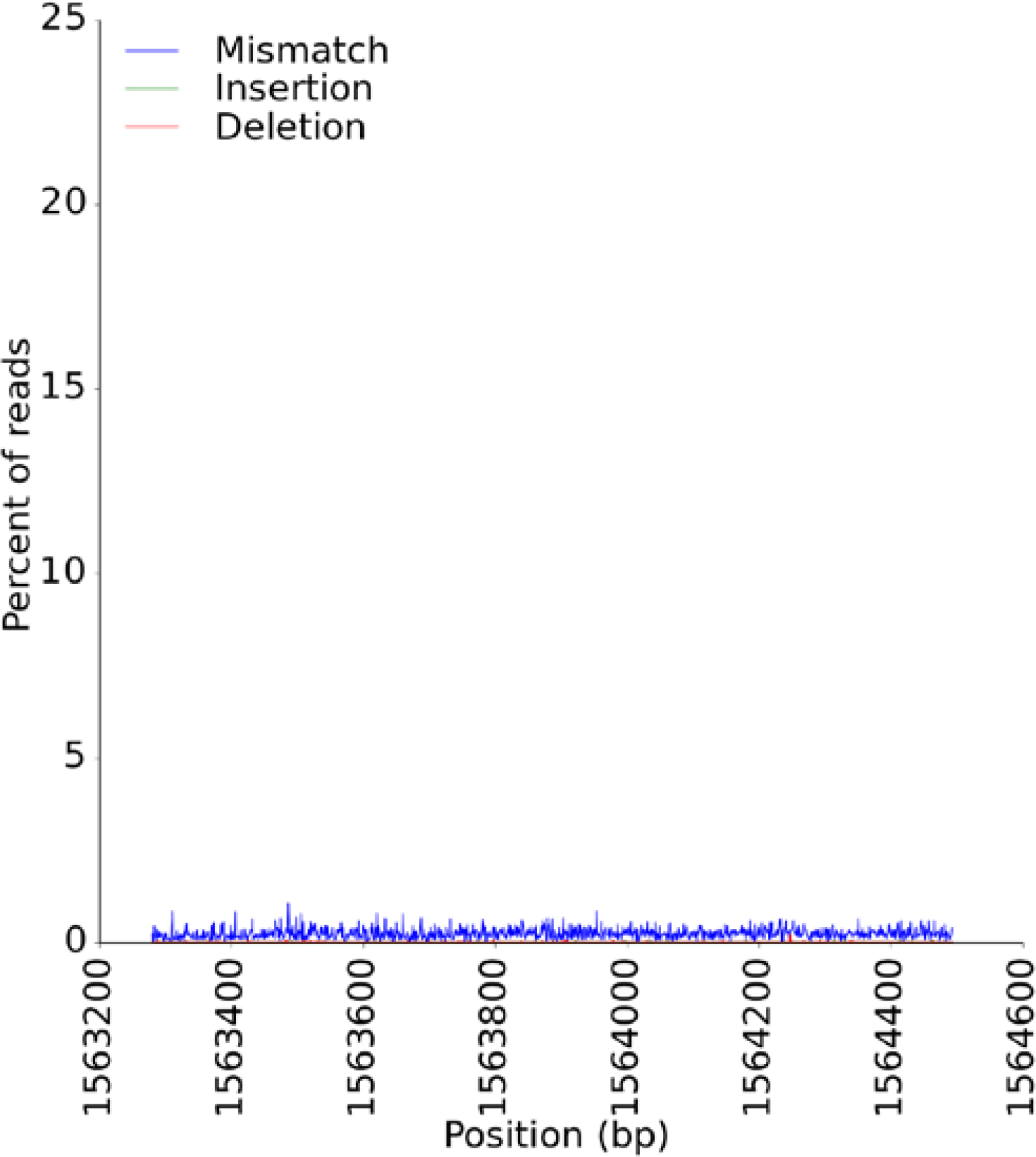
Read-level MiSeq mismatch and insertion/deletion calls in the coding region (+/-200bp) of the RGATCY-targeting *M.CsaI* gene in *C. salexigens*.

**Supplementary Figure 15:**
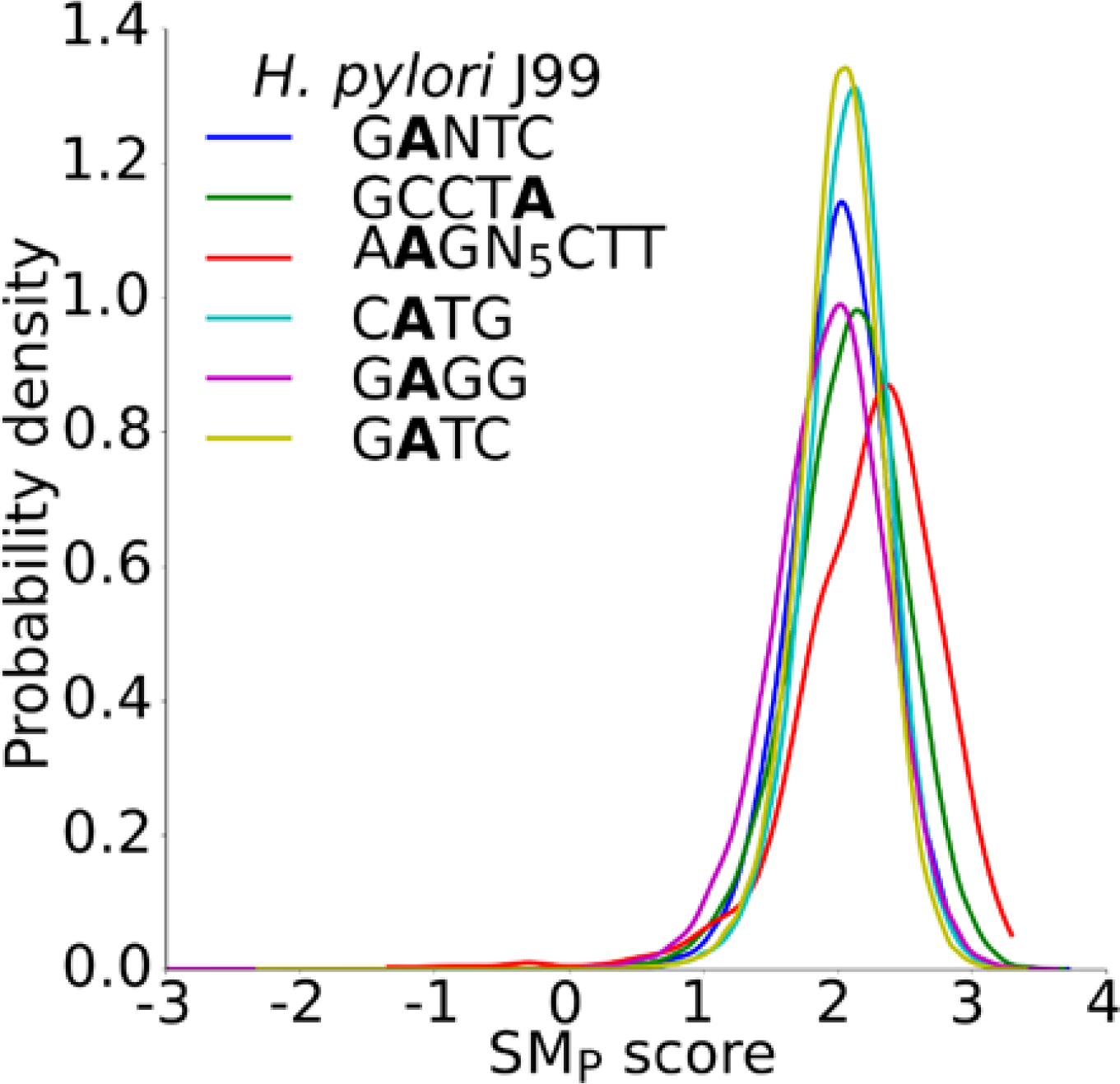
SM_p_ score distributions (at least 10 motif sites per molecule) for all 6mA motifs in *H. pylori* J99 (excluding GWCAY, TCAN6TRG, and TCNNGA) that contained at least ten distinct motif sites on at least 500 molecules.

**Supplementary Figure 16:**
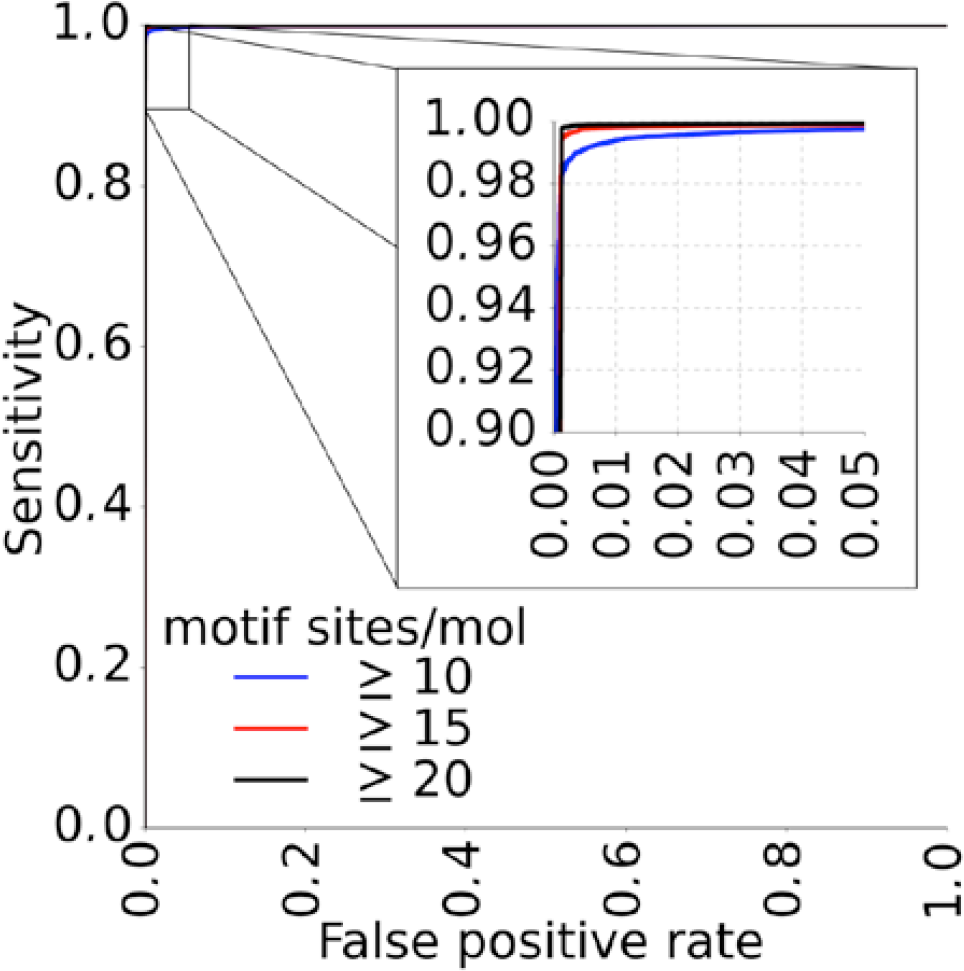
Sensitivity and specificity of the SM_p_ method for detecting molecules that are methylated at the GATC motif in *H. pylori* J99.

**Supplementary Figure 17:**
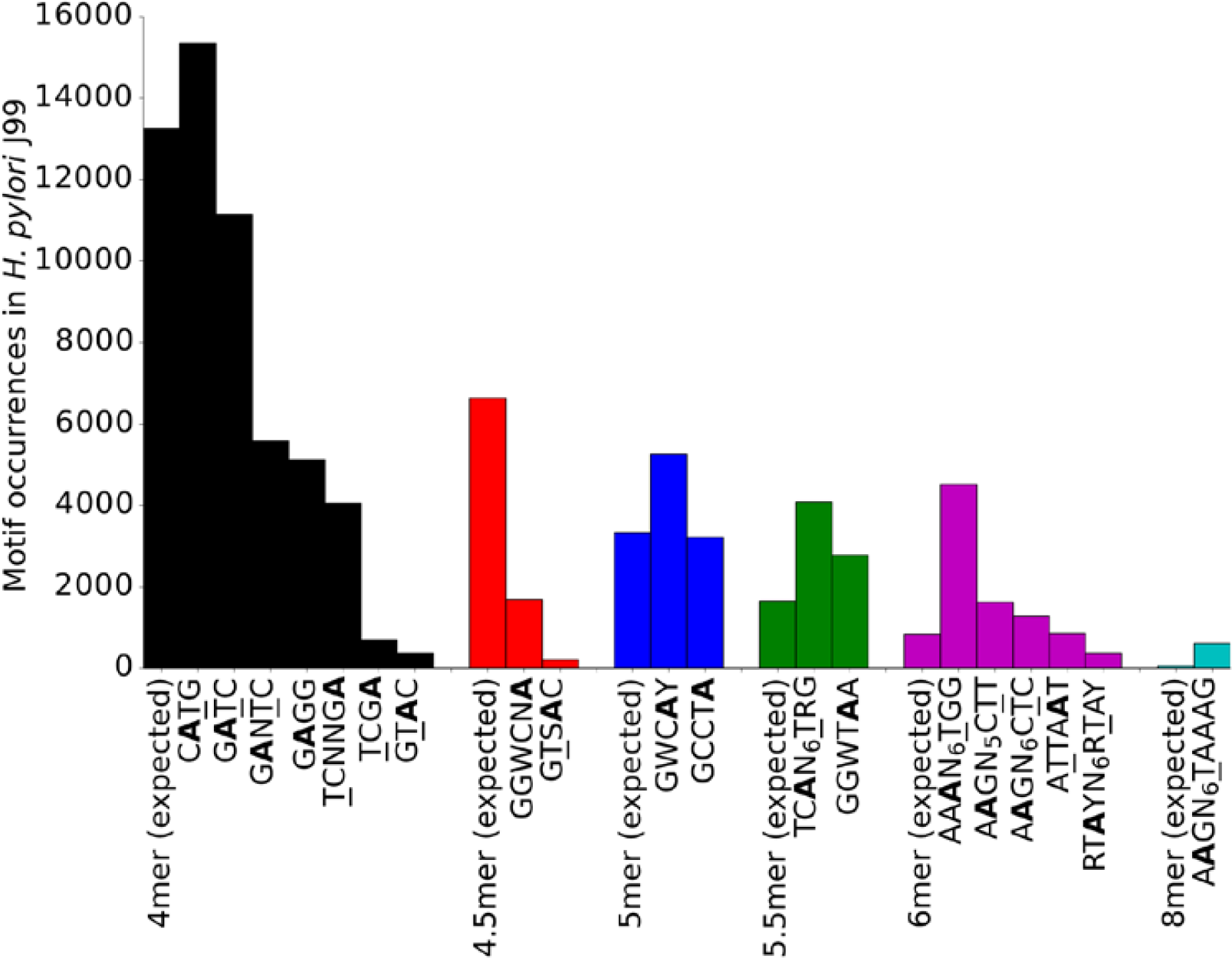
Count of expected vs. observed motif occurrences in the *H. pylori* J99 genome based on k-mer size.

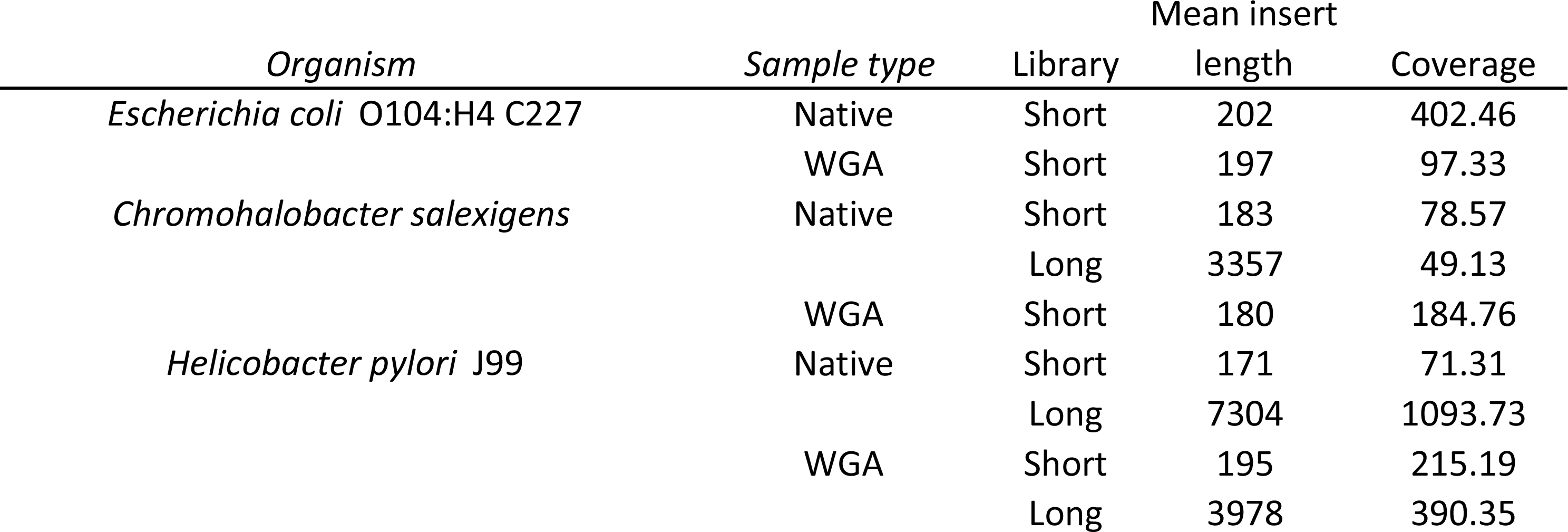

| Mean read<br>length | Mean subread<br>length |
| --- | --- |
| 5609 | 172 |
| 6091 | 167 |
| 8119 | 183 |
| 5233 | 3357 |
| 9023 | 180 |
| 4735 | 168 |
| 7965 | 6871 |
| 6873 | 167 |
| 4957 | 3978 |

